# A Unified Molecular Signature of Systemic Lupus Erythematosus Revealed by Integrated, Multi-Cohort Transcriptomic Analysis

**DOI:** 10.1101/834093

**Authors:** Winston A. Haynes, D. James Haddon, Vivian K. Diep, Avani Khatri, Erika Bongen, Gloria Yiu, Imelda Balboni, Christopher R. Bolen, Rong Mao, Paul J. Utz, Purvesh Khatri

**Affiliations:** Institute for Immunity, Transplantation and Infection, Stanford University School of Medicine, Stanford, CA, USA; Department of Medicine, Division of Biomedical Informatics Research, Stanford University School of Medicine, Stanford, CA, USA; Department of Medicine, Division of Immunology and Rheumatology, Stanford University School of Medicine, Stanford, CA, USA; Department of Pediatrics, Division of Allergy, Immunology and Rheumatology, Stanford University School of Medicine, Stanford, CA, USA

## Abstract

Systemic lupus erythematosus (SLE) is a complex autoimmune disease that follows an unpredictable disease course and affects multiple organs and tissues. We performed an integrated, multi-cohort analysis of 7,471 transcriptomic profiles from 40 independent studies to identify robust gene expression changes associated with SLE. We identified a 93-gene signature (SLE MetaSignature) that is differentially expressed in the blood of SLE patients compared to healthy volunteers; distinguishes SLE from other autoimmune, inflammatory, and infectious diseases; and persists across diverse tissues and cell types. The SLE MetaSignature correlated significantly with disease activity and other clinical measures of inflammation. We prospectively validated the SLE MetaSignature in an independent cohort of pediatric SLE patients using a microfluidic RT-qPCR array. We found that 14 of the 93 genes in the SLE MetaSignature were independent of interferon-induced and neutrophil-related transcriptional profiles that have previously been associated with SLE. Pathway analysis revealed dysregulation associated with nucleic acid biosynthesis and immunometabolism in SLE. We further refined a neutropoeisis signature and identified under-appreciated transcripts related to immune cells and oxidative stress. Our multi-cohort, transcriptomic analysis has uncovered under-appreciated genes and pathways associated with SLE pathogenesis, with the potential to advance clinical diagnosis, biomarker development, and targeted therapeutics for SLE.

## Introduction

Systemic lupus erythematosus (SLE) is a complex, heterogeneous, chronic autoimmune disease that can affect multiple organs and tissues, including the skin, kidneys, joints, lungs, blood, and central nervous system. SLE follows an unpredictable disease course, punctuated by periods of flare and remission(1). High titer, class-switched antibodies that bind to nuclear antigens including dsDNA, ribonucleoprotein (RNP), Smith, SSA (Ro), and SSB (La), are used in the diagnosis and monitoring of SLE, and are thought to be pathogenic. The heterogeneity of SLE makes it challenging for clinicians to manage. Identification of robust molecular changes associated with SLE, despite the patient heterogeneity, will likely improve our understanding and management of SLE.

A number of gene expression studies have shed light on the molecular pathogenesis of SLE. For example, microarray analyses of blood cells derived from SLE patients have shown that the interferon (IFN) pathway is dysregulated in a subset of individuals who have more active and severe disease(2–5). Increases in IFN-related genes have also been observed in subsets of patients with other diseases, including systemic sclerosis (SSc), dermatomyositis (DM), polymyositis (PM), primary Sjögren’s syndrome (SS), and rheumatoid arthritis (RA), although levels of IFN-inducible gene products were typically highest in SLE(6–10). A review of the biomedical literature identified IFN and neutrophils as a major focus of recent SLE research, with approximately 150 and 40 references per year, respectively. In addition to the IFN signature, up-regulation of transcripts associated with granulopoiesis and plasmablasts were observed in individuals who have SLE and found to be associated with disease activity(3, 5). McKinney *et al.* used gene expression analysis of purified immune cell populations to identify a transcriptional signature in CD8 T cells that was associated with increased likelihood of SLE disease flare(11). They went on to identify an exhaustion signature, associated with decreased risk of flare, in CD8 T cells from individuals who have SLE(12). However, the majority of these studies have been limited by small sample sizes, low levels of clinical and geographic heterogeneity, potential artifacts related to use of a single experimental gene array platform, and lack of external validation. A more robust approach is needed to interrogate the molecular signatures that underlie the highly variable presentation and course of SLE.

We have previously described a multi-cohort analysis framework (MetaIntegrator) to identify robust disease signatures, and repeatedly demonstrated its applications for discovering novel diagnostics, prognostics and drug targets, and drug repurposing, which leverages the biological and technical heterogeneity present in the large amounts of publicly-available gene expression data across a broad spectrum of conditions including infections, organ transplant, vaccination, cancer, and autoimmune diseases(13–15). MetaIntegrator is based on a random-effects meta-analysis, drawing statistical power from the integration of many diverse datasets(14). By computing effect sizes for each dataset independently, MetaIntegrator embraces heterogeneity and avoids the limitations of batch effect correction. We have demonstrated application of this framework across a broad spectrum of diseases including cancer(16, 17), solid organ transplant(18), sepsis(19), viral infection(20), tuberculosis(21), neurodegenerative diseases(22), vaccination(23), and systemic sclerosis(24). Here, we applied the framework to analyze 40 publicly-available whole transcriptome profile datasets containing 7,471 samples from SLE patients, individuals with other autoimmune diseases or infections, and healthy volunteers. Together, these datasets represented real-world diversity because of both (i) the biological heterogeneity, as the samples were collected from multiple tissue and cell types (e.g. blood, skin, and kidney) at 17 centers across five countries; and (ii) the technical heterogeneity, as data were generated using diverse microarray platforms (e.g. Affymetrix arrays, Illumina beadchips, and Hitachisoft chips). Our analysis identified a robust SLE MetaSignature that (i) distinguishes SLE from other autoimmune and inflammatory diseases; (ii) is present in multiple affected tissues and immune cell subsets; (iii) is independent of age; and (iv) is correlated with disease activity. We validated the SLE MetaSignature using additional independent publicly-available transcript datasets. We then devised a custom, microfluidic RT-qPCR assay to analyze RNA transcripts in blood derived from a prospective, independent pediatric SLE (pSLE) cohort. Pathway analysis identified novel dysregulated pathways in SLE, including those related to nucleotide biosynthesis and metabolism. Importantly, we identified a non-IFN component of the SLE MetaSignature that correlated more positively with disease activity measures than the IFN-related genes. Finally, our results discovered 14 “non-IFN, non-neutrophil” genes as under-appreciated targets for biomarker and therapeutic development.

## Results

### Identification of the SLE MetaSignature

To perform a comprehensive, unbiased study of the molecular changes underlying SLE, we identified and downloaded gene expression data from all publicly-available human SLE datasets in Gene Expression Omnibus(25). In total, we identified 40 datasets from 17 centers in 5 countries comprised of 7,471 samples derived from whole blood, PBMCs, kidney, skin, synovium, B cells, T cells, monocytes, neutrophils, and endothelial progenitor cells [Figure 1, Table 1]. We randomly selected 6 datasets consisting of 370 whole blood and PBMC samples as “Discovery” datasets, based on our previous finding that 5 datasets with 250-300 samples are sufficient to find a robust disease gene signature using our multi-cohort analysis framework(26). We divided the remaining 34 datasets into “Validation” (2,407 samples in 8 datasets) and “Extended Validation” datasets (4,694 samples in 26 datasets). Discovery and Validation datasets were required to include PBMC or whole blood samples from healthy controls and SLE patients. Extended Validation datasets included samples from other tissues or cell types, comparisons between SLE and other diseases, and longitudinal SLE samples.

**Figure 1.**
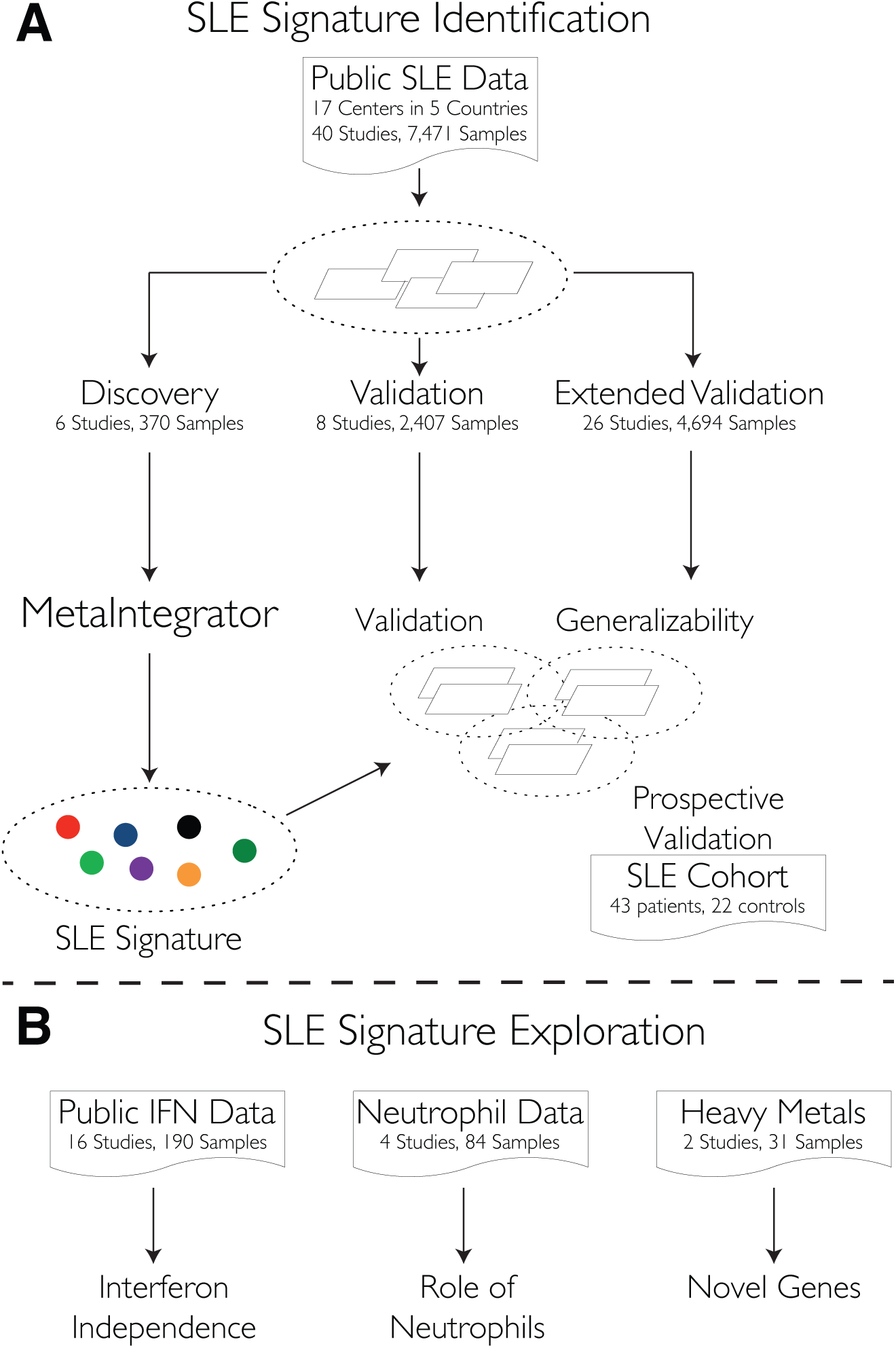
Identification and validation of a SLE-specific gene signature using integrated, multi-cohort analysis. (A) We downloaded 40 publicly-available datasets from 17 centers in five countries comprising 7,471 samples. We identified datasets that included whole blood or PBMC samples from SLE patients and healthy volunteers to serve as discovery (6 studies) and validation (8 studies) sets. The remaining 26 studies contained samples from other tissue types or lacked healthy volunteer samples, and were examined as extended validation datasets. We used the MetaIntegrator framework to identify a 93 gene SLE MetaSignature (effect size > 1, FDR < .05, measured in ≥ 4 datasets). We examined the classification accuracy of the signature in validation data and the generalizability of the signature in the extended validation data. To prospectively validate the SLE meta-analysis signature using an external cohort, we analyzed individuals who have pSLE (n = 43) or JIA (n = 12) from the Stanford Pediatric Rheumatology Clinic, as well as healthy adult (n = 10) volunteers using Fluidigm RT-qPCR arrays. (B) We leveraged publicly-available data to identify non-IFN components of the SLE MetaSignature, examine the role of neutrophils in SLE, and study heavy metal exposure. pSLE: pediatric SLE.

**Table 1.**
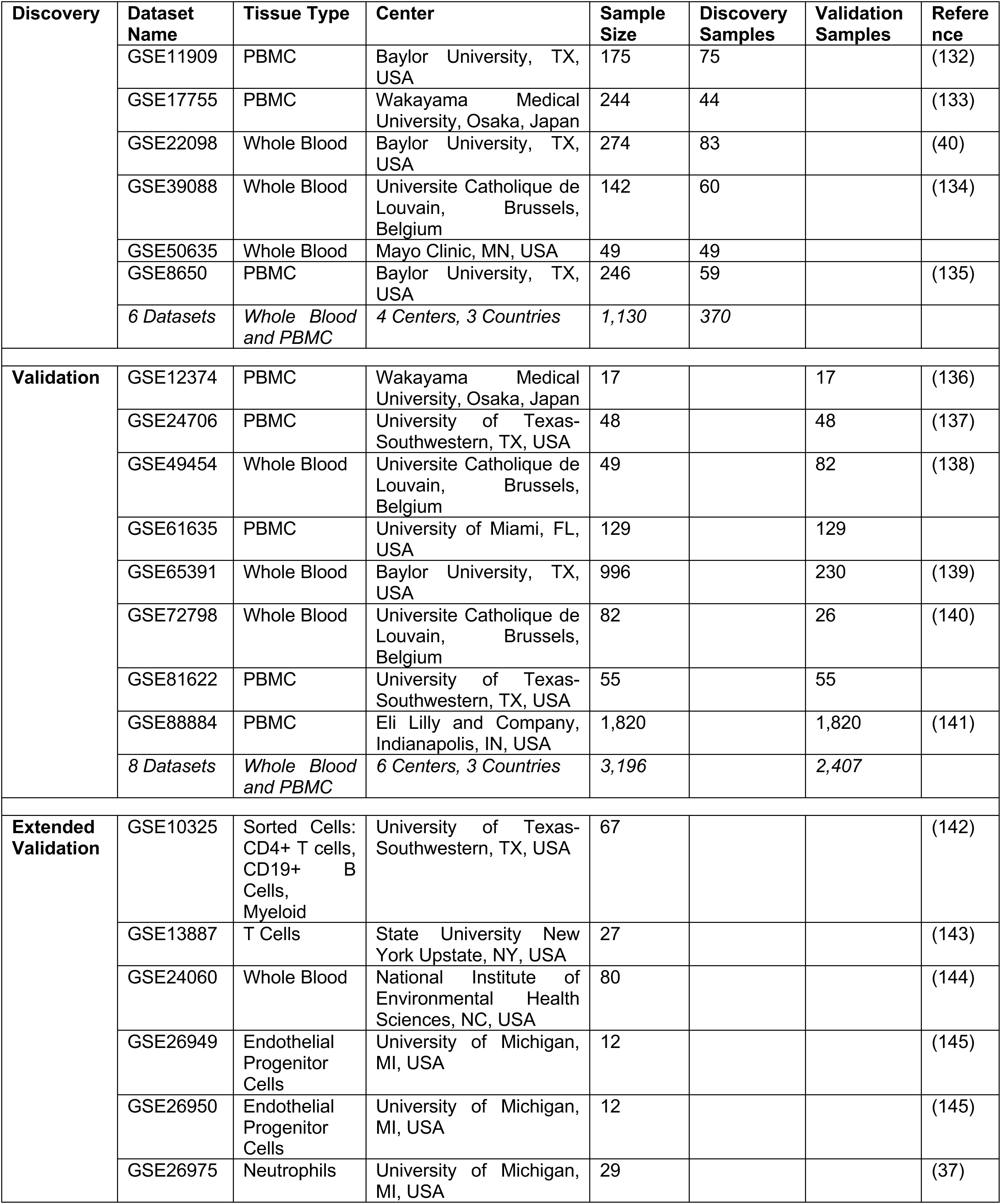
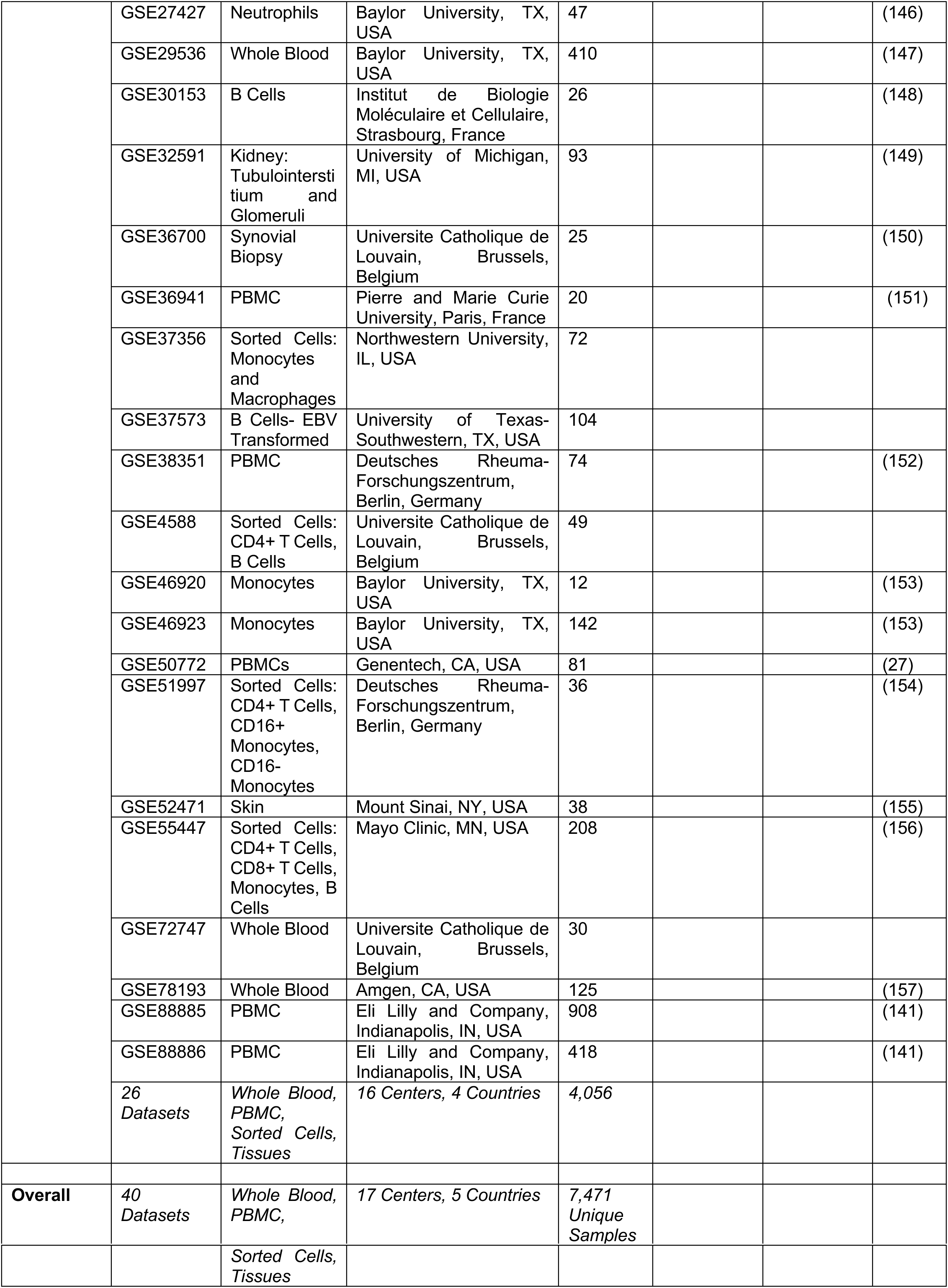
SLE dataset summaries. For every study included in this paper, we summarized the number of patients, tissue of origin, study location, and associated publications. More extensive descriptions are provided in Supplement S1.

| Discovery | Dataset Name | Tissue Type | Center | Sample Size | Discovery Samples | Validation Samples | Reference |
| --- | --- | --- | --- | --- | --- | --- | --- |
|  | GSE11909 | PBMC | Baylor University, TX, USA | 175 | 75 |  | (132) |
|  | GSE17755 | PBMC | Wakayama Medical University, Osaka, Japan | 244 | 44 |  | (133) |
|  | GSE22098 | Whole Blood | Baylor University, TX, USA | 274 | 83 |  | (40) |
|  | GSE39088 | Whole Blood | Universite Catholique de Louvain, Brussels, Belgium | 142 | 60 |  | (134) |
|  | GSE50635 | Whole Blood | Mayo Clinic, MN, USA | 49 | 49 |  |  |
|  | GSE8650 | PBMC | Baylor University, TX, USA | 246 | 59 |  | (135) |
|  | 6 Datasets | Whole Blood and PBMC | 4 Centers, 3 Countries | 1,130 | 370 |  |  |
| Validation | GSE12374 | PBMC | Wakayama Medical University, Osaka, Japan | 17 |  | 17 | (136) |
|  | GSE24706 | PBMC | University of Texas-Southwestern, TX, USA | 48 |  | 48 | (137) |
|  | GSE49454 | Whole Blood | Universite Catholique de Louvain, Brussels, Belgium | 49 |  | 82 | (138) |
|  | GSE61635 | PBMC | University of Miami, FL, USA | 129 |  | 129 |  |
|  | GSE65391 | Whole Blood | Baylor University, TX, USA | 996 |  | 230 | (139) |
|  | GSE72798 | Whole Blood | Universite Catholique de Louvain, Brussels, Belgium | 82 |  | 26 | (140) |
|  | GSE81622 | PBMC | University of Texas-Southwestern, TX, USA | 55 |  | 55 |  |
|  | GSE88884 | PBMC | Eli Lilly and Company, Indianapolis, IN, USA | 1,820 |  | 1,820 | (141) |
|  | 8 Datasets | Whole Blood and PBMC | 6 Centers, 3 Countries | 3,196 |  | 2,407 |  |
| Extended Validation | GSE10325 | Sorted Cells: CD4+ T cells, CD19+ B Cells, Myeloid | University of Texas-Southwestern, TX, USA | 67 |  |  | (142) |
|  | GSE13887 | T Cells | State University New York Upstate, NY, USA | 27 |  |  | (143) |
|  | GSE24060 | Whole Blood | National Institute of Environmental Health Sciences, NC, USA | 80 |  |  | (144) |
|  | GSE26949 | Endothelial Progenitor Cells | University of Michigan, MI, USA | 12 |  |  | (145) |
|  | GSE26950 | Endothelial Progenitor Cells | University of Michigan, MI, USA | 12 |  |  | (145) |
|  | GSE26975 | Neutrophils | University of Michigan, MI, USA | 29 |  |  | (37) |
|  | GSE27427 | Neutrophils | Baylor University, TX, USA | 47 |  |  | (146) |
|  | GSE29536 | Whole Blood | Baylor University, TX, USA | 410 |  |  | (147) |
|  | GSE30153 | B Cells | Institut de Biologie Moléculaire et Cellulaire, Strasbourg, France | 26 |  |  | (148) |
|  | GSE32591 | Kidney: Tubulointerstitium and Glomeruli | University of Michigan, MI, USA | 93 |  |  | (149) |
|  | GSE36700 | Synovial Biopsy | Universite Catholique de Louvain, Brussels, Belgium | 25 |  |  | (150) |
|  | GSE36941 | PBMC | Pierre and Marie Curie University, Paris, France | 20 |  |  | (151) |
|  | GSE37356 | Sorted Cells: Monocytes and Macrophages | Northwestern University, IL, USA | 72 |  |  |  |
|  | GSE37573 | B Cells- EBV Transformed | University of Texas-Southwestern, TX, USA | 104 |  |  |  |
|  | GSE38351 | PBMC | Deutsches Rheuma-Forschungszentrum, Berlin, Germany | 74 |  |  | (152) |
|  | GSE4588 | Sorted Cells: CD4+ T Cells, B Cells | Universite Catholique de Louvain, Brussels, Belgium | 49 |  |  |  |
|  | GSE46920 | Monocytes | Baylor University, TX, USA | 12 |  |  | (153) |
|  | GSE46923 | Monocytes | Baylor University, TX, USA | 142 |  |  | (153) |
|  | GSE50772 | PBMCs | Genentech, CA, USA | 81 |  |  | (27) |
|  | GSE51997 | Sorted Cells: CD4+ T Cells, CD16+ Monocytes, CD16- Monocytes | Deutsches Rheuma-Forschungszentrum, Berlin, Germany | 36 |  |  | (154) |
|  | GSE52471 | Skin | Mount Sinai, NY, USA | 38 |  |  | (155) |
|  | GSE55447 | Sorted Cells: CD4+ T Cells, CD8+ T Cells, Monocytes, B Cells | Mayo Clinic, MN, USA | 208 |  |  | (156) |
|  | GSE72747 | Whole Blood | Universite Catholique de Louvain, Brussels, Belgium | 30 |  |  |  |
|  | GSE78193 | Whole Blood | Amgen, CA, USA | 125 |  |  | (157) |
|  | GSE88885 | PBMC | Eli Lilly and Company, Indianapolis, IN, USA | 908 |  |  | (141) |
|  | GSE88886 | PBMC | Eli Lilly and Company, Indianapolis, IN, USA | 418 |  |  | (141) |
|  | 26 Datasets | Whole Blood, PBMC, Sorted Cells, Tissues | 16 Centers, 4 Countries | 4,056 |  |  |  |
| **Overall** | 40 Datasets | Whole Blood, PBMC, | 17 Centers, 5 Countries | 7,471 Unique Samples |  |  |  |
|  |  | *Sorted Cells,*<br>*Tissues* |  |  |  |  |  |

We identified 93 significantly differentially regulated genes (82 up-regulated and 11 down-regulated; Table S1) with a false discovery rate (FDR) ≤ 5% and an absolute effect size ≥ 1 compared to healthy volunteers in the Discovery datasets [Figure 2A, Table S1]. We defined these 93 genes as the “SLE MetaSignature.” In the Validation datasets, 73 of these 93 SLE MetaSignature genes met the same filtering criteria (|ES| ≥ 1 and FDR ≤ 5%) and effect sizes for all 93 genes exhibited the same directionality as in the Discovery datasets [Figure 2B, Figure S1]. Of the 20 SLE MetaSignature genes that did not meet the filtering criteria, 18 were statistically significant (FDR ≤ 5%), but had an effect size less than 1 (median effect size 0.78). but had an effect size less than 1 (median effect size 0.78). In the Extended Validation datasets, which included data from diverse sample types and other diseases, the SLE MetaSignature gene effect sizes were consistent with the Discovery dataset [Figure 2C]. Regardless of the genetic background of the patients, technical variation, tissue, and cell type, the genes comprising the SLE MetaSignature were all differentially expressed [Figure 2A-C], demonstrating the robustness of the SLE MetaSignature.

**Figure 2.**
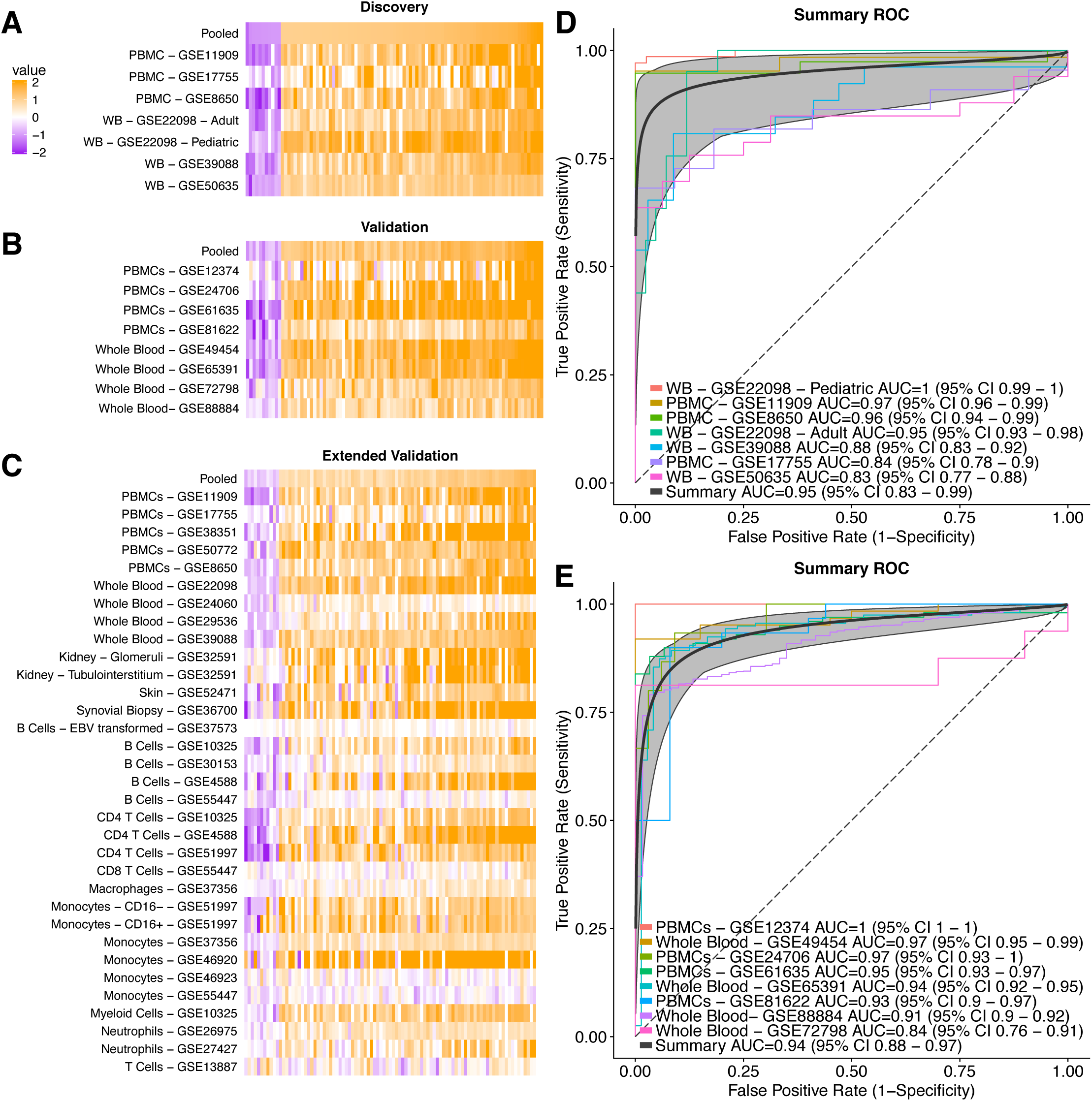
SLE MetaSignature persists across diverse datasets. Effect size heatmaps of SLE MetaSignature genes across (A) discovery, (B) validation, and (C) extended validation datasets. Each column represents a gene in the SLE MetaSignature, ordered from lowest- to highest-effect size in the discovery data. Each row represents a gene expression dataset. Receiver operating characteristic curves are broken into (D) discovery and (E) validation data. A perfect classifier will have an AUROC of 1 and a random classifier will have an AUROC of 0.5. We show both whole blood (WB) and peripheral blood mononuclear cell (PBMCs) samples. The summary curve is a composite of the individual study curves. The extended validation ROC plot is shown in Figure S9.

We defined an “SLE MetaScore” for each sample using the 93-gene signature (see Methods). In the Discovery datasets, the SLE MetaScore distinguished SLE patient samples from healthy samples with a summary area under the receiver operating characteristic curve (AUROC) of 0.95 (95% confidence interval (CI) 0.83-0.99) [Figure 2D]. The SLE MetaScore distinguished samples from SLE patients and healthy volunteers with high accuracy in the 8 Validation datasets (summary AUROC=0.94, 95% CI = 0.89-0.97) [Figure 2E], further demonstrating the robustness of the SLE MetaSignature.

Of the 93 genes in the SLE MetaSignature, 46 had been previously associated with SLE(2, 3, 5, 27). To the best of our knowledge, the remaining 47 genes have not previously been associated with SLE. We performed pathway analysis of the SLE MetaSignature using Differential Expression Analysis for Pathways (DEAP)(28) to identify biological processes that are dysregulated in SLE. DEAP takes advantage of the meta-analysis effect sizes for all genes (not just those in the SLE MetaSignature) and pathway topology to identify patterns of differential expression that are consistent with known biological pathways. By taking advantage of effect sizes of all genes, DEAP significantly improves power compared to gene list based approaches(28). Further, DEAP specifies genes involved in the most differentially expressed subpathway. As input for DEAP, we used study level effect sizes from the Discovery and Validation datasets(28). Table 2 summarizes pathways that were differentially expressed at a false discovery rate (FDR) ≤ 0.10 based on 5,000 random rotations of the data. In addition to the expected inflammatory pathways (e.g. IFNγ signaling pathway, chemokine/cytokine-mediated signaling pathway, and interleukin signaling pathway), our analysis identified several highly significant, unexpected pathways (salvage pyrimidine deoxyribonucleotides, formyltetrahydrofolate biosynthesis, and salvage pyrimidine ribonucleotides) related to nucleic acid metabolism. Thus, pathway analysis of the SLE MetaSignature provided insights into the biological mechanisms underlying SLE.

**Table 2.**
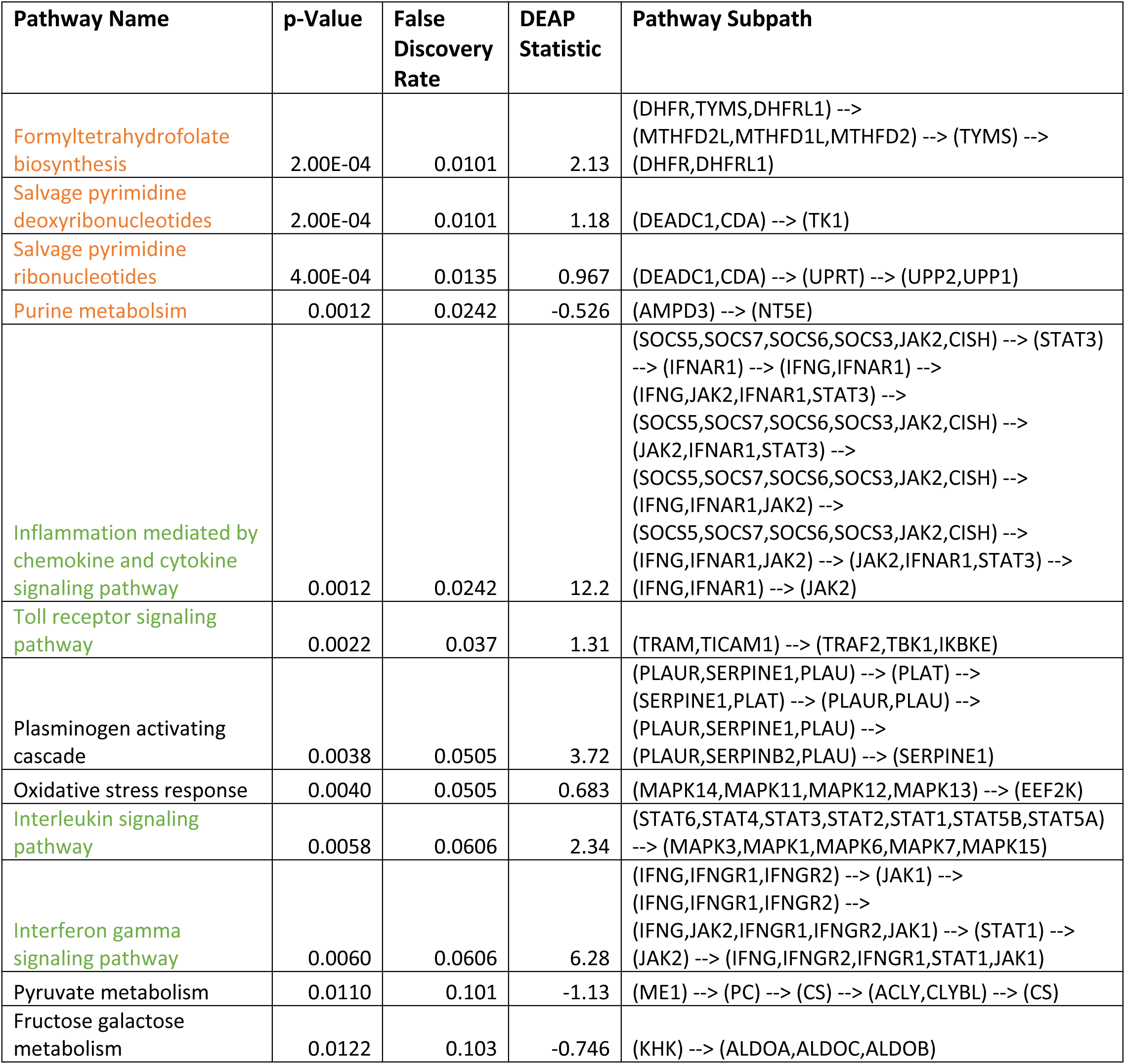
Results from pathway analysis of the SLE meta-analysis data. . Pathways related to inflammation (green) and nucleic acid metabolism (orange).

### SLE MetaScore distinguishes SLE from other autoimmune, inflammatory, and infectious diseases

We compared SLE MetaScores across inflammatory conditions, including other autoimmune and infectious diseases to explore its specificity to SLE. We found that adult SLE (aSLE) and pediatric SLE (pSLE) patients had significantly higher SLE MetaScores than individuals with staphylococcal infection, streptococcal pharyngitis, Still’s disease (systemic onset juvenile idiopathic arthritis, sJIA), RA, pyogenic pyoderma gangrenosum and acne (PAPA), B cell deficiency, diabetes, human immunodeficiency virus infection, and liver transplant acute rejection in whole blood and PBMC samples across multiple independent datasets [Figure 3A-B and Figure S2A-B]. In concordance with the previously reported increased severity of disease observed in pediatric SLE patients compared to adults(29), we found that pediatric SLE patients had significantly higher SLE MetaScores compared to adult SLE patients [Figure 3B]. Taken together, these results demonstrate that, both in adult and pediatric populations, the SLE MetaScore is highly specific to SLE compared to other autoimmune, inflammatory, and infectious diseases.

**Figure 3.**
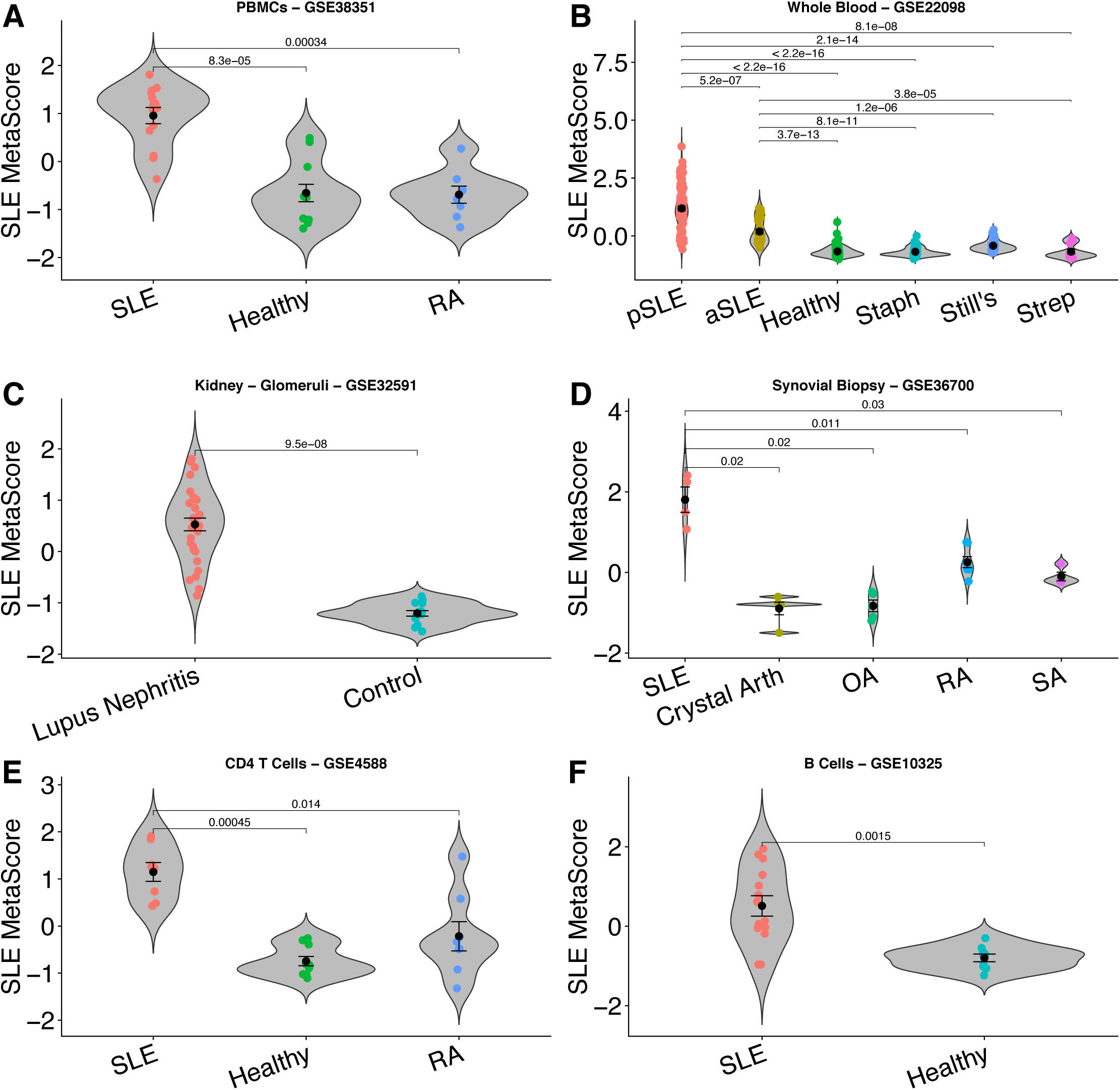
SLE MetaSignature persists across diseases, tissues, and cell types. In these violin plots, each point represents a patient and the SLE MetaScore (y-axis) has been calculated using the SLE MetaSignature genes. (A-B) The SLE MetaScore distinguished SLE from other diseases. See also Figure S2. (C-D The SLE MetaScore distinguishes SLE from other diseases and healthy controls in diverse tissues. See also Figure S3. (E-F) The SLE MetaScore distinguishes SLE patients from healthy and other diseases in sorted immune cells. See also Figure S4. For all panels: rheumatoid arthritis (RA); pediatric SLE (pSLE); adult SLE (aSLE); staphylococcal infection (Staph); Still’s disease (Still’s); streptococcal pharyngitis (Strep); microcrystalline arthritis (Crystal Arth); osteoarthritis (OA); and seronegative arthritis (SA).

### The SLE MetaScore is systemically higher across tissues in SLE patients

SLE is a systemic autoimmune disease that affects multiple tissues and organs. Therefore, we explored whether the SLE MetaScore is persistent in tissues other than whole blood and PBMCs in SLE patients. SLE MetaScores were higher in a dataset derived from glomeruli and tubulointerstitium of kidneys from individuals with SLE compared with pre-transplant living donors [Figure 3C and Figure S3A]. SLE MetaScores were higher in a dataset from synovial biopsies of SLE patients compared to those with microcrystalline arthritis (gout and pseudogout), osteoarthritis (OA), RA, or seronegative arthritis [Figure 3D]. Finally, we found that a dataset derived from skin biopsies from individuals with discoid lupus erythematosus exhibited significantly higher SLE MetaScores than healthy volunteers and individuals with psoriasis, suggesting shared pathways between systemic and cutaneous lupus [Figure S3B]. Collectively, these results provide strong evidence that the SLE MetaScore is higher in multiple affected tissues in SLE in comparison to both healthy controls and other autoimmune diseases.

### The SLE MetaScore is differentially expressed in diverse immune cell types

Multiple functional changes have been described in T cells of SLE patients, including upregulation of co-stimulatory molecules, hypomethylation, increased expression of key immune-related genes(30), and aberrant signaling pathway activation downstream of TCR activation(31). We found that the SLE MetaScore was significantly higher in multiple independent datasets from CD4+ T cells of SLE patients compared to healthy volunteers [Figures S4A-C] and RA patients [Figure 3E]. Similarly, the SLE MetaScore was significantly increased in a dataset from CD8+ T cells of individuals with SLE, compared to healthy volunteers [Figure S4D].

Dysregulation of B cells is a hallmark of SLE, including autoantibody production, defective negative selection, and changes in the proportions of key B cell subpopulations(32, 33). The SLE MetaScore was less robust in datasets from B cells than T cells, classifying SLE in some datasets [Figure 3F and Figure S4E] but not others [Figure S4F-G]. Finally, the SLE MetaScores in datasets from monocytes and neutrophils were not significantly different between SLE patients and healthy controls [data not shown].

### The SLE MetaScore is positively correlated with disease activity and inflammation

The SLE Disease Activity Index (SLEDAI) is a standardized, albeit imperfect, measure of disease severity and activity. SLEDAI is based on the presence or absence of 24 features at the time of the visit, including arthritis, rash, fever, and increases in anti-DNA auto-antibodies. It is often used by clinicians to monitor disease activity in an individual SLE patient(3). Five independent datasets that profiled PBMC or whole blood samples from SLE patients also reported SLEDAI scores. We observed a positive correlation between SLEDAI and the SLE MetaScore across each of the five datasets [Figure 4A-B, Figure S5A-D]. The median correlation across these studies (0.281) was significantly elevated compared to random gene sets (p < 0.01). The weakest SLEDAI correlation is observed in GSE27427, which contains only 18 samples and is derived from neutrophils. The positive correlation of SLEDAI with SLE MetaScore in the blood is notable, considering the SLE MetaSignature was identified without considering disease activity when selecting initial datasets for discovery. Further, we found that the SLE MetaScore correlated highly with individual clinical measures of systemic inflammation including erythrocyte sedimentation rate (ESR) [Figure S5E](34), and levels of complement C3 [Figure 4C] and C4 [Figure S5F].

**Figure 4.**
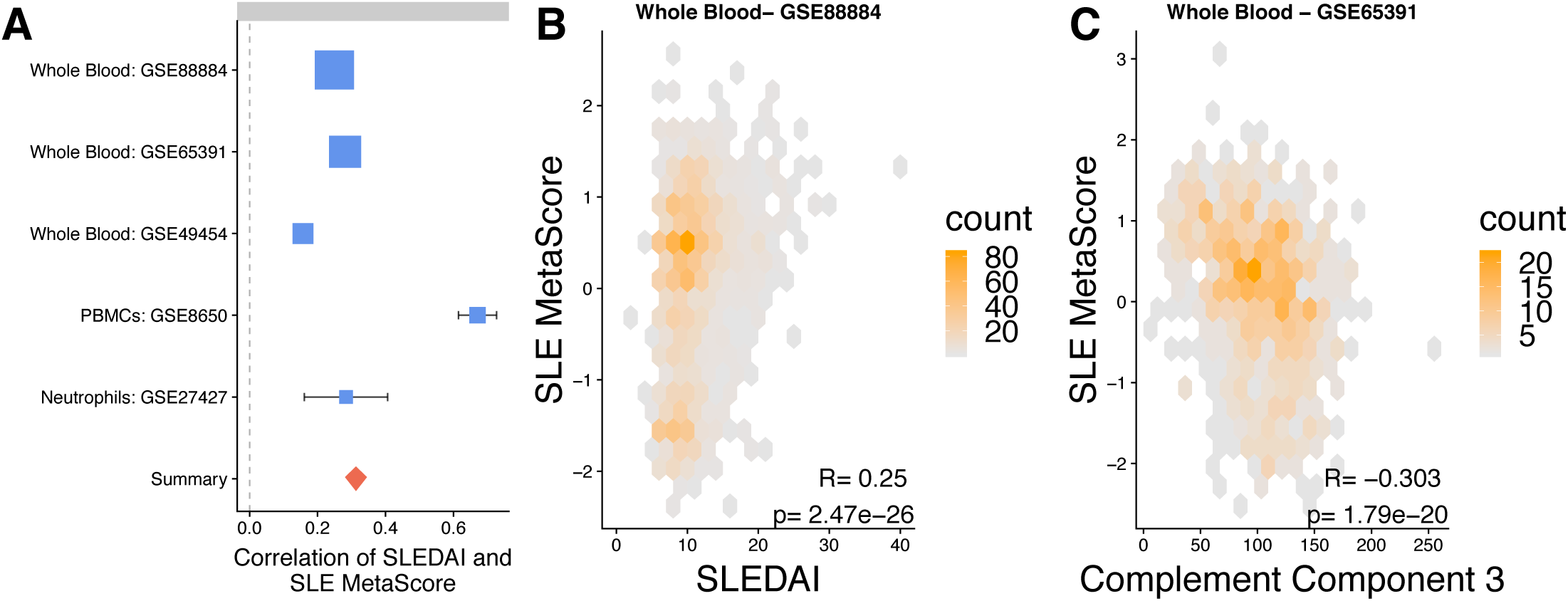
Disease activity is significantly associated with the SLE MetaScore. (A) Positive spearman correlations are observed across all five datasets where SLEDAI was available. Box size is proportional to the confidence of the correlation estimate. Summary is the pooled, inverse variance summary correlation value. (B) The SLE MetaScore in an example SLE whole blood datasets is positively correlated with SLEDAI. (C) SLE MetaScore is inversely correlated with complement C3 levels. In all panels, every point represents a single patient. See also Figure S5.

### Prospective validation of SLE MetaSignature in an independent pSLE cohort

We validated the SLE MetaSignature in an independent pediatric cohort by studying RNA transcripts in whole blood samples from healthy adult controls, and pediatric patients with SLE or JIA. We selected 33 genes from the SLE MetaSignature based on their significance and availability of validated probes for measuring expression using a microfluidic RT-qPCR array [Table S4]. Thirty genes out of 33 were significantly differentially expressed in SLE samples compared to healthy adult controls and pediatric JIA patients [FDR < 5%, Table S4]. Further, the SLE MetaScores based on these 33 genes in the pediatric SLE patients were significantly higher than healthy adult controls and pediatric JIA patients [p=3.7e-5 and 1.8e-6, respectively; Figure 5A]; distinguished pediatric SLE patients with high accuracy (AUROC=0.94); and were positively correlated with SLEDAI [Spearman correlation=0.307, p=0.045; Figure 5B].

**Figure 5.**
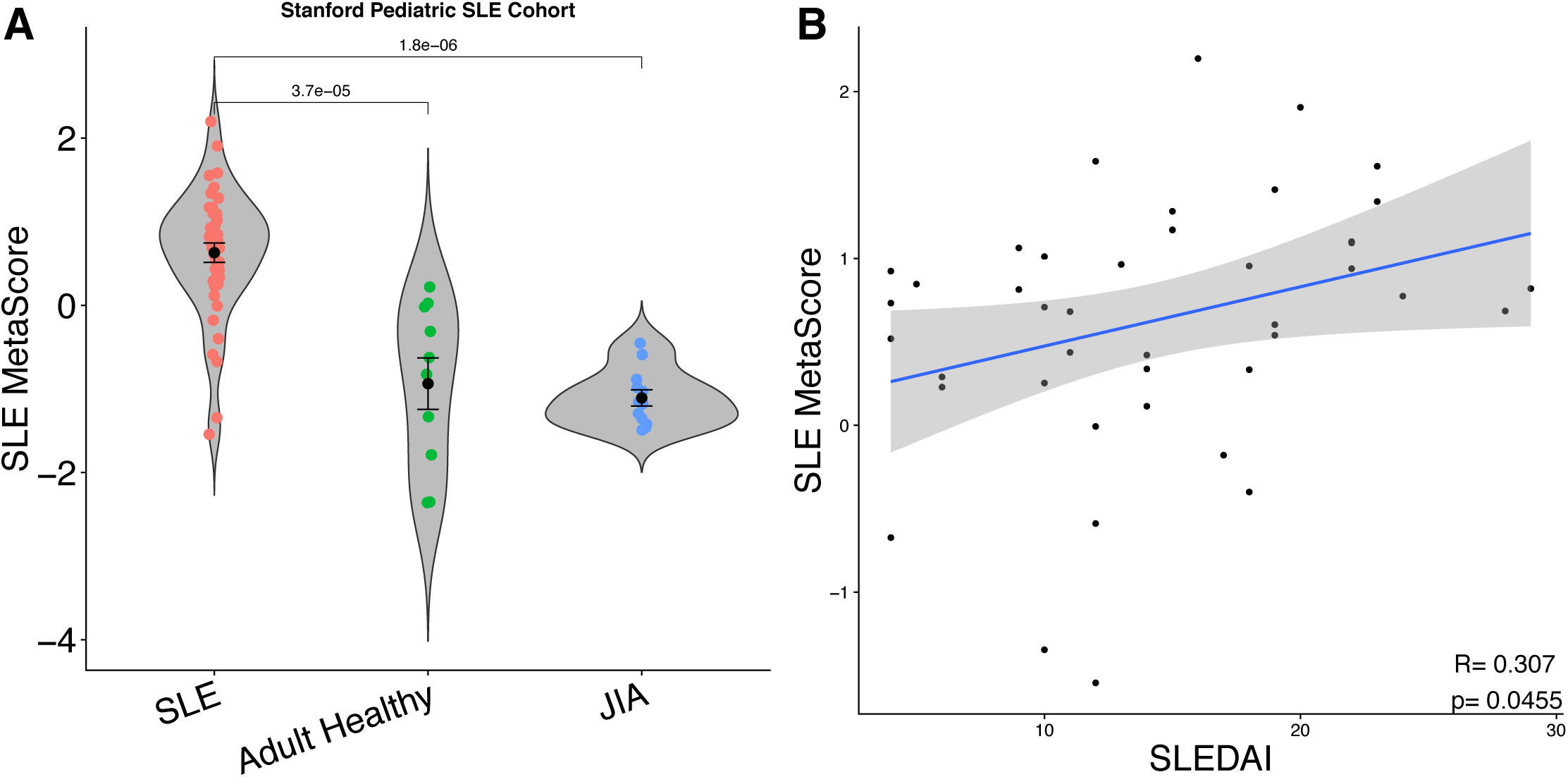
Prospective validation of the SLE MetaSignature in an independent pSLE cohort using a microfluidic RT-qPCR assay. The relative levels of 33 transcripts selected from the SLE MetaSignature (and housekeeping genes) were analyzed in total RNA prepared from the whole blood of new onset pSLE patients (n=43), individuals with JIA (n=12), and adult healthy volunteers (n=10) in parallel using a multiplexed, microfluidic RT-qPCR assay. (A) The geometric means of the relative concentrations of the 33 transcripts in the SLE MetaSignature were calculated for each individual. Plots show Z-scores calculated across individuals. Mann-Whitney tests were used to compare groups. (B) SLEDAI scores for individual patients were calculated at the time of sampling and were correlated with their SLE MetaScores.

### A subset of the SLE MetaSignature is not robustly induced by IFN

Dysregulation of the type I IFN pathway has been repeatedly observed in subsets of SLE patients with active disease, and is thought to be a critical mediator in disease pathology. Therefore, we explored the proportion of IFN-stimulated genes in the SLE MetaSignature. We analyzed 16 transcriptome datasets composed of 190 samples derived from primary human cells treated with type I IFN to identify a robust set of type I IFN-stimulated genes [Table S2]. Of the 93 genes in the SLE MetaSignature, 70 were significantly differentially expressed (effect size > 0.8) in primary cells stimulated by type I IFN [Figure 6A, Table S1]. The remaining 23 genes in the SLE MetaSignature had low effect sizes and high FDRs within the IFN-stimulated datasets [Figure S6], suggesting that these 23 genes were not affected in cells exposed to type I IFN.

**Figure 6.**
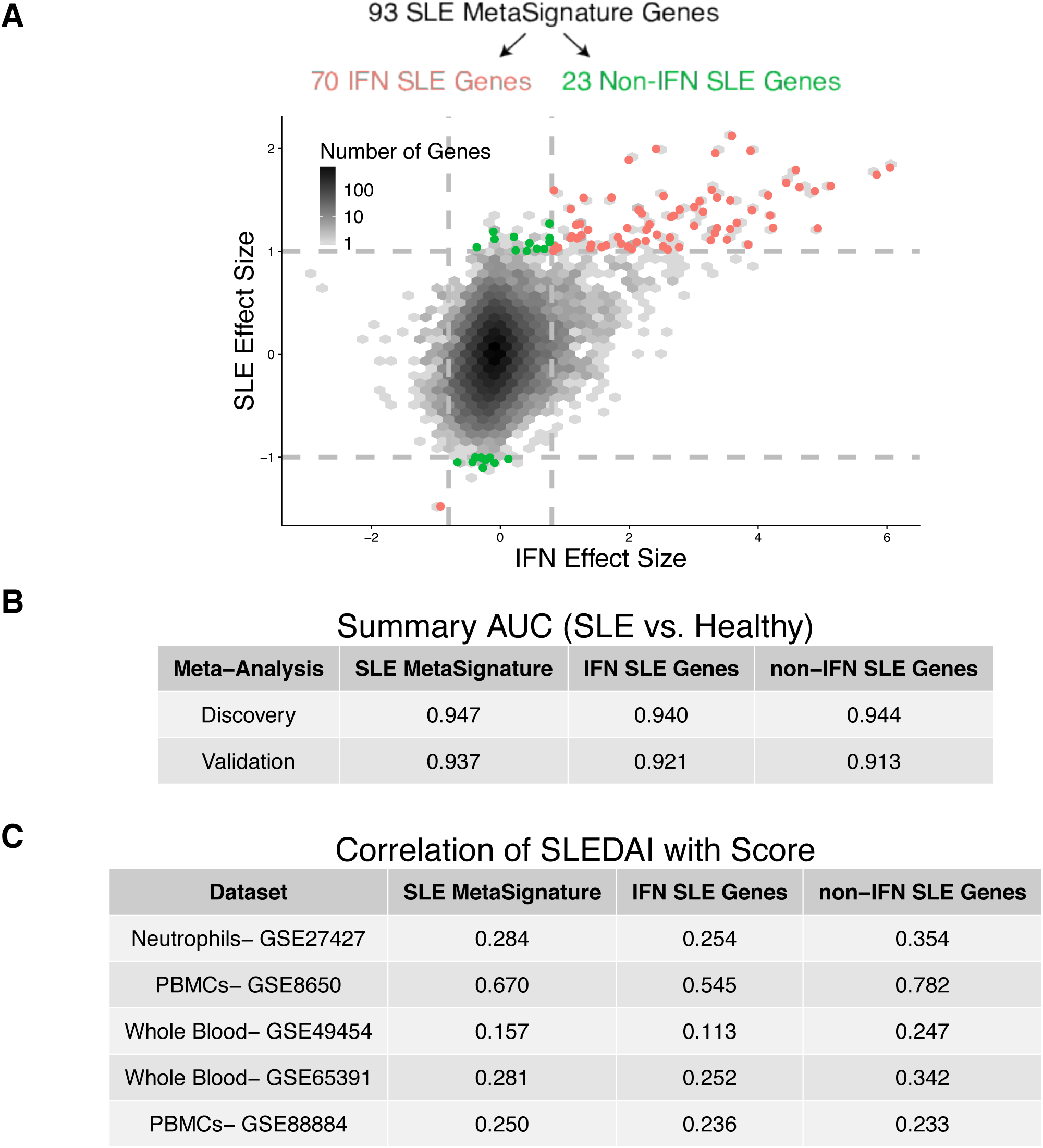
SLE MetaSignature genes dependent or independent of IFN stimulation. Based on our meta-analysis of 16 datasets of type I IFN stimulation in primary cells, we estimated IFN effect sizes in response to stimulation. We compared the SLE MetaSignature to the results of an IFN meta-analysis by examining (A) SLE effect size vs. IFN effect size. For a volcano plot for all of the IFN effect sizes, please see [Figure S6]. We compared the complete SLE MetaSignature, IFN SLE MetaSiganture, and non-IFN SLE MetaSignature in terms of (B) summary AUC in discovery and validation and (C) correlation with SLEDAI in 5 datasets.

We separated the SLE MetaSignature into “IFN” and “non-IFN” SLE MetaSignatures, and computed score as before. Both scores distinguished SLE patients with equally high accuracy in the validation datasets [Figure 6B]. In four out of five datasets with SLEDAI disease severity measurements, the non-IFN SLE MetaSignature had a higher correlation with SLEDAI than the IFN SLE MetaSignature [Figure 6C]. Collectively, our analyses identified a clinically important, non-IFN component of the SLE MetaSignature.

### The role of non-IFN MetaSignature genes in neutrophils

We used immunoStates to identify cell lineages that most highly express genes that comprise the SLE MetaSignature. We found that many of the non-IFN SLE MetaSignature genes were up-regulated in neutrophils(35), consistent with prior literature implicating neutrophils in SLE (3, 36–40). Low density granulocytes exhibit enhanced type I IFN production and NETosis, a form of neutrophil cell death implicated in SLE pathogenesis(41) in which DNA neutrophil extracellular traps (NETs) are extruded from activated neutrophils(38, 41). We identified a transcript profiling dataset that compared low density granulocytes and neutrophils from SLE patients or healthy controls. We observed that the non-IFN SLE MetaSignature was prominently found in low density granulocytes from SLE patients, but not in neutrophils from SLE patients or healthy controls(37) [Figure 7A]. We observed a strong correlation between neutrophil abundance and SLE MetaScore in both studies where quantitative neutrophil counts were available [Figure S10]. Collectively, these results suggest that the SLE MetaSignature genes related to neutrophils are the result of an expansion of the neutrophil compartment in SLE patients rather than an altered expression profile in SLE neutrophils.

**Figure 7.**
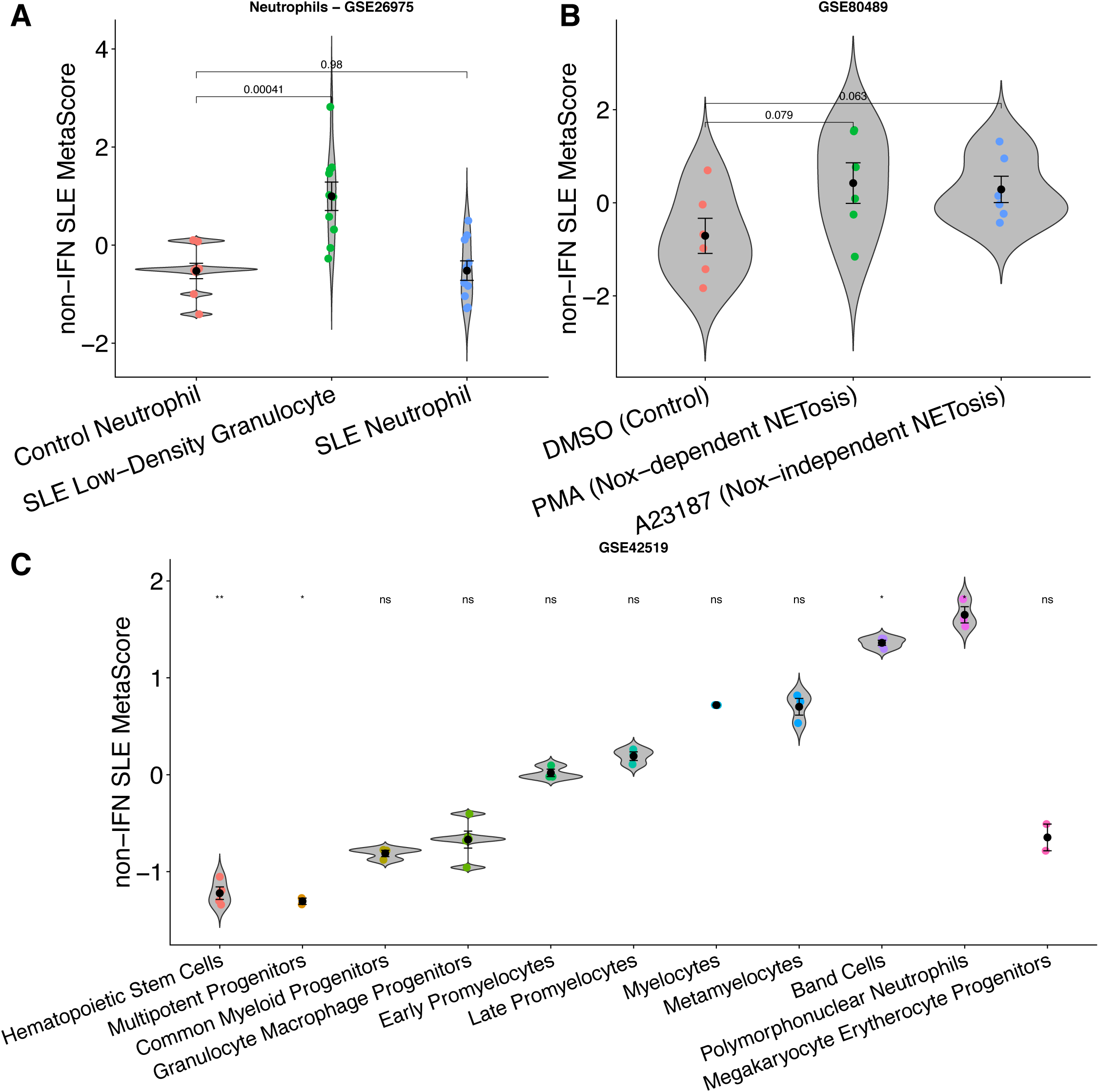
The role of non-IFN MetaSignature genes in SLE neutrophils. We examined the non-IFN component of the SLE MetaSignature (a total of 23 genes) in datasets related to neutrophils. (A) Non-IFN SLE MetaScore in control neutrophils, SLE neutrophils, and SLE low-density granulocytes (LDGs). SLE MetaScore in LDGs is significantly elevated compared to both the control neutrophil and SLE neutrophil populations. (B) Non-IFN SLE MetaScore enriched in primary cells in response to both Nox-dependent (PMA) and Nox-independent (A23187) NETosis. (C) Non-IFN SLE MetaScore progressively increased in sorted intermediate cell populations along the neutropoeisis lineage. See also Figure S7.

To further explore the role of the non-IFN genes in neutrophils, we identified four publicly-available gene expression datasets with 84 samples that explored either NETosis or neutrophil development [Table S3]. The non-IFN SLE MetaSignature was up-regulated in cell lines that were stimulated to induce both Nox-dependent and Nox-independent NETosis [Figure 7B]. The non-IFN SLE MetaSignature progressively increases during intermediate stages of neutropoeisis [Figure 7C and Figure S7A-B]. Collectively, these results indicate that a significant proportion of the non-IFN SLE MetaSignature is related to transcriptional signatures of NETosis and neutropoeisis.

### Identification of “under-appreciated, non-IFN, non-neutrophil” SLE MetaSignature genes

IFN stimulation and neutrophil involvement explained the differential expression of 79 of the 93 genes in the SLE MetaSignature [Figure 8A, Table S1]. The remaining 14 genes (termed “Under-appreciated SLE MetaSignature”, [Table 3]) provided an opportunity to explore new disease mechanisms that underlie SLE. The Under-appreciated SLE MetaScore correlated more positively with disease activity measurements than the IFN SLE MetaScore in every blood derived dataset [Table S5]. Interestingly, three members of the metallothionein family (*MT1E*, *MT1F*, and *MT1HL1*) were in the Under-appreciated SLE MetaSignature. Metallothioneins play an important role in oxidative stress responses and the clearance of heavy metals. We identified two datasets in which human cell lines were exposed chronically to cadmium or acutely to zinc. The Under-appreciated SLE MetaSignature was significantly elevated in cells exposed to heavy metals when compared to the untreated cell lines(42) [Figure 8B-C], providing a potential link between SLE and heavy metals, or with other environmental stimuli that induce oxidative stress. A cadre of the remaining 11 genes in the Under-appreciated SLE MetaSignature encode molecules with interesting functions related to immune cells, while the remainder of the genes have not been linked to SLE and have yet to be well characterized in the literature.

**Table 3.**
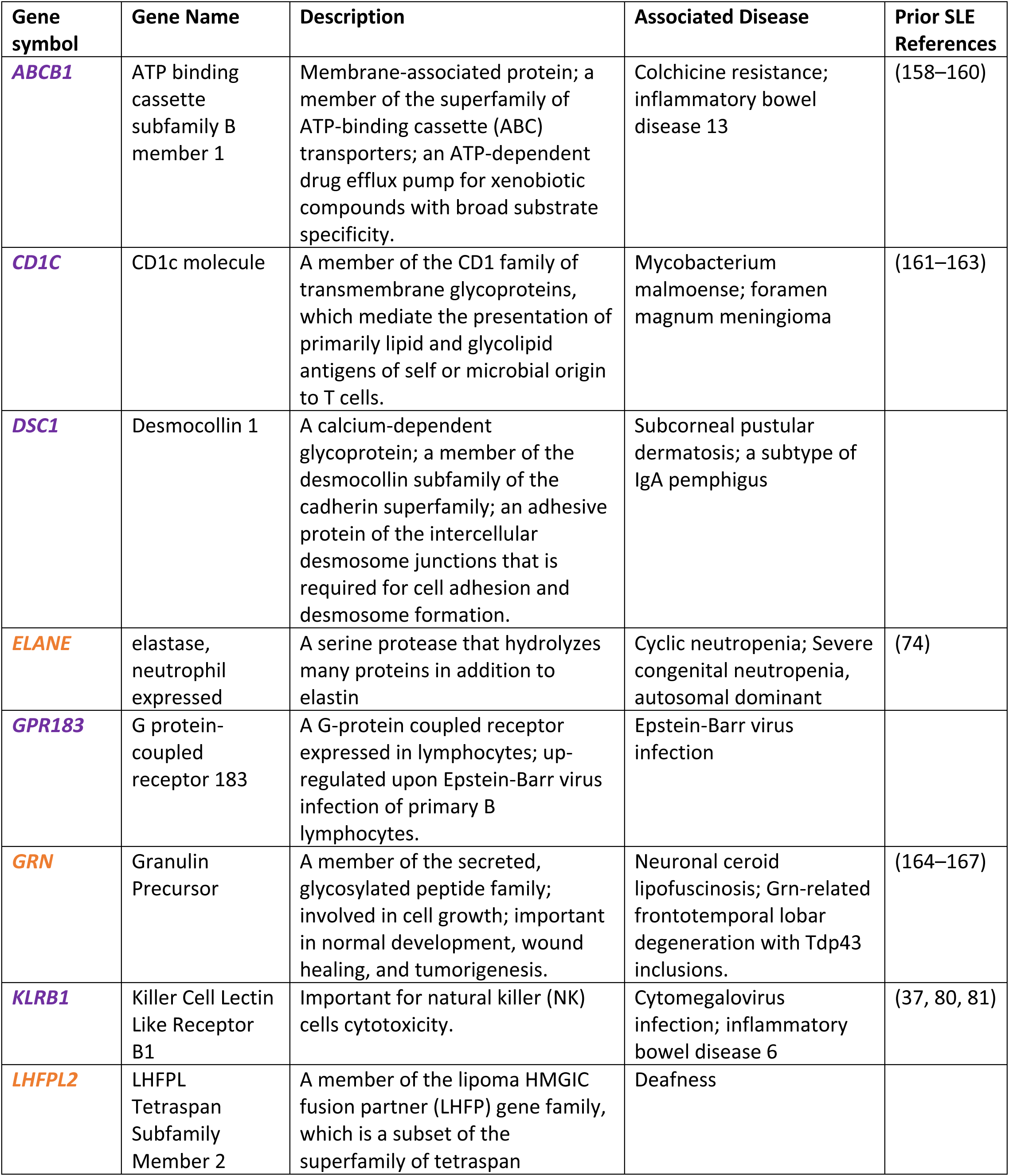
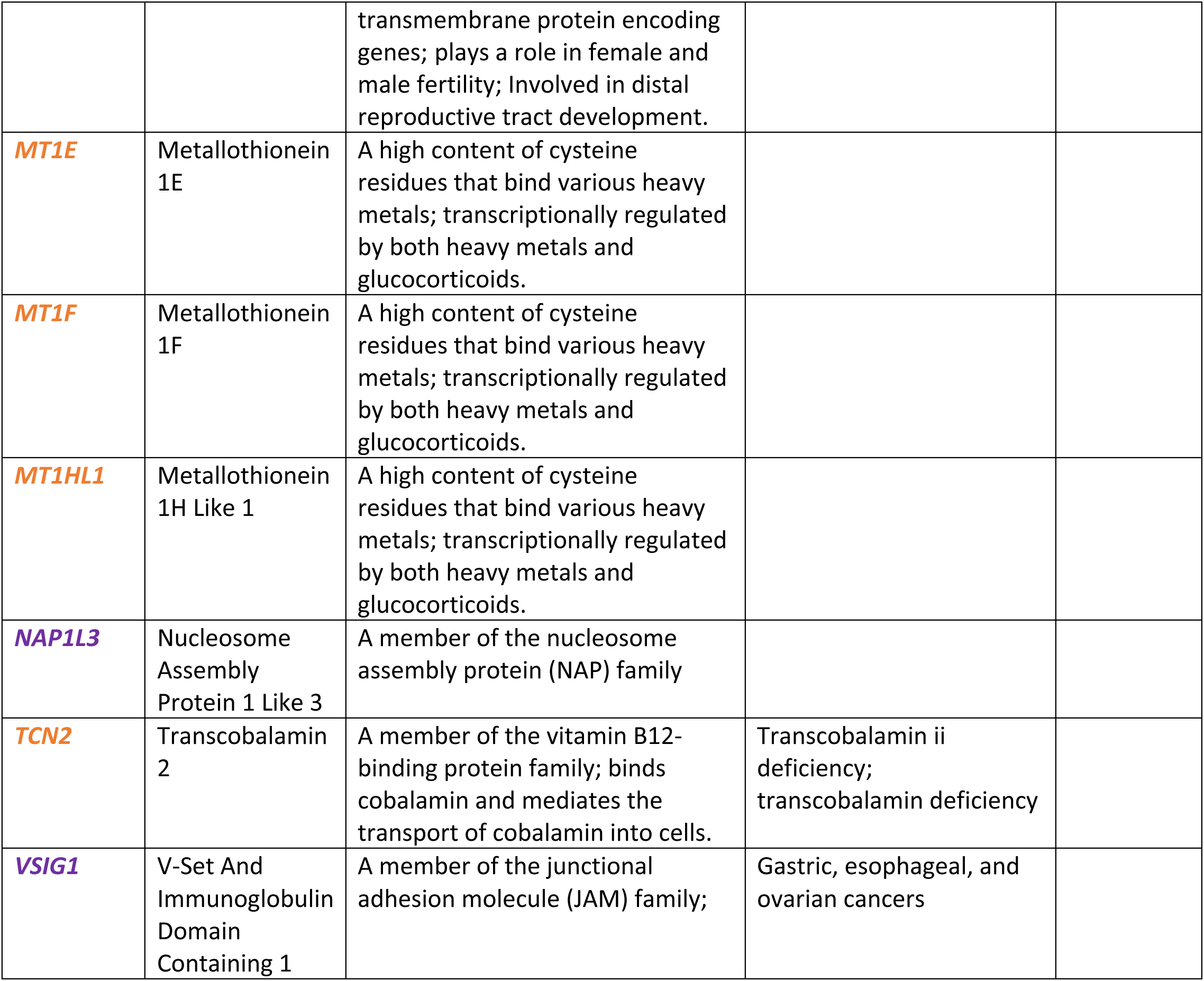
Under-appreciated genes in SLE MetaSignature. The 14 genes from the Under-appreciated SLE MetaSignature are listed, including their gene symbol, gene name, description, and human diseases with which these genes have associated. Genes are colored by whether they are up (orange) or down (purple) regulated in SLE vs. healthy controls.

**Figure 8.**
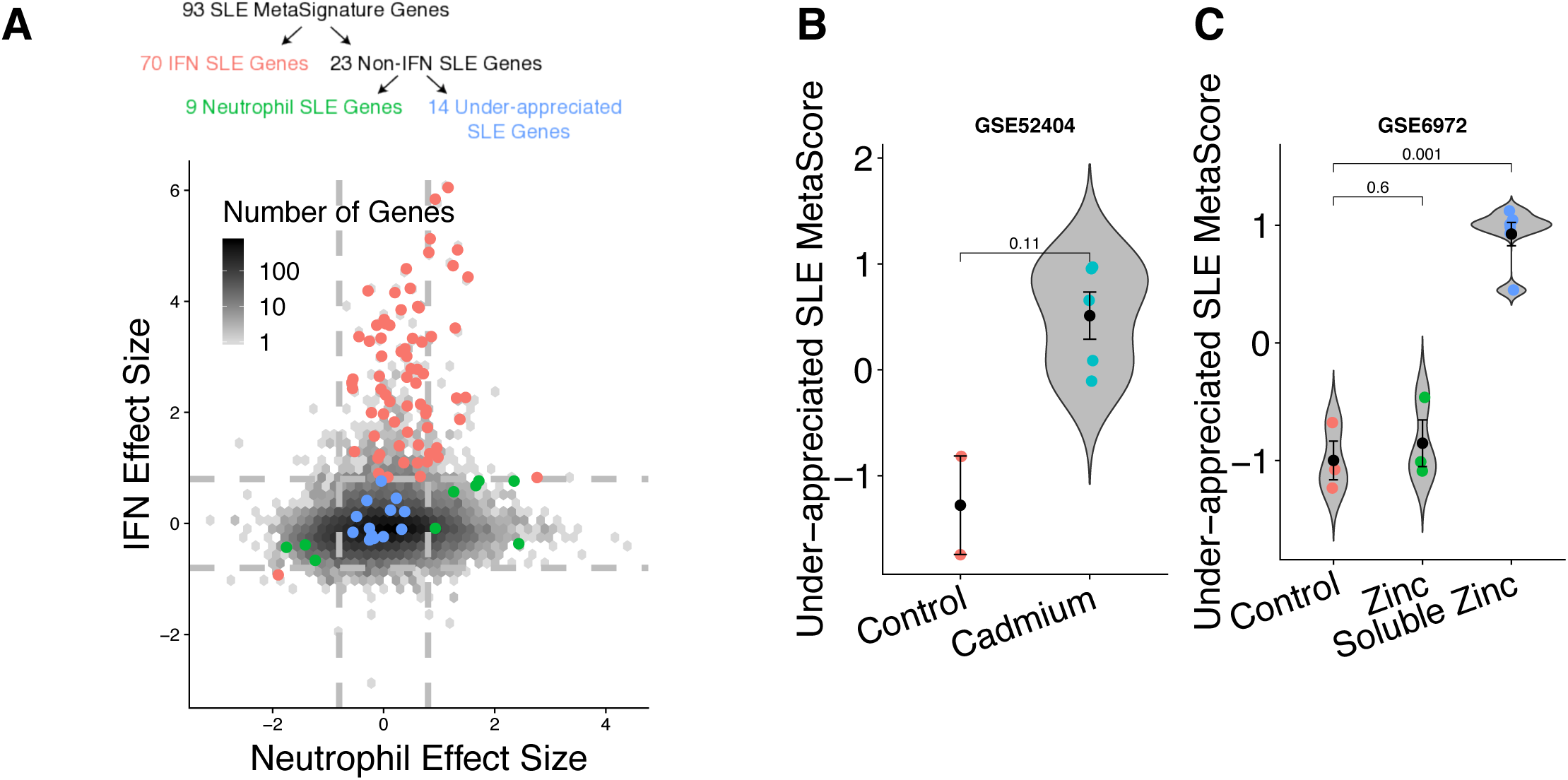
Identification of under-appreciated non-IFN, non-neutrophil SLE MetaSignature genes. (A) IFN effect size vs. neutrophil effect size. Neutrophil effect size estimated from immnoStates(35). Red indicates the 70 genes which were in the SLE MetaSignature and were significantly differentially expressed in response to IFN. Green indicates the 9 genes which were in the SLE MetaSignature, were not significantly differentially expressed in response to IFN, and were significantly differentially expressed in neutrophils. Blue indicates the 14 genes in the SLE MetaSignature that were not significantly differentially expressed in neutrophils or in response to IFN stimulation. Dashed lines indicate an effect size threshold of |0.8| for both neutrophil and IFN effect sizes. (B) Cell lines which were chronically exposed to cadmium displayed an increased Under-appreciated SLE MetaScore compared to control cell lines. (C) Cells exposed to a water soluble zinc compound exhibited an increased Under-appreciated SLE MetaScore compared to those exposed to both a control compound and an insoluble form of zinc(42).

## Discussion

Previous gene expression meta-analyses in SLE have been limited to few experiments, lacked external validation, or did not investigate the signature’s specificity to SLE(43, 44). Our method leverages biological and technical heterogeneity to identify a robust disease signature, and has been successful in diverse diseases that range from cancer to autoimmunity and infection(16–24). We performed a multi-cohort gene expression analysis of over 7,000 samples from 40 datasets representing real-world biological heterogeneity (including genetic background, age, sex, treatment, tissue, cell type, and disease duration) and technical heterogeneity (including RNA isolation, microarray platform, sample preparation, and experimental protocol) to identify a persistent SLE MetaSignature. The robustness and reproducibility of the SLE MetaSignature demonstrate its generalizability to diverse patient populations not observed in traditional, single-cohort analyses(14).

Beyond generalizability, the SLE MetaSignature was both specific to SLE and correlated with disease activity. Since the SLE MetaSignature distinguished SLE from other diseases such as diffuse or organ specific autoimmune diseases, inflammatory arthritides, and infectious diseases, the SLE MetaSignature identified SLE-specific disease processes instead of those that are generically dysregulated in other immune-mediated diseases. SLEDAI is the current standard for assessing severity of SLE disease activity although it is a qualitative, subjective, and difficult-to-reproduce measure(45). Thus, the positive correlation between the SLE MetaScore and SLEDAI suggests that the SLE MetaScore is capturing disease activity, but is quantitative and objective. Therefore, it could potentially serve as a metric of disease activity in future studies, or an exploratory outcome measure in future clinical trials. Because the SLE MetaScore includes both IFN and non-IFN genes, it expands upon the current best practices of using IFN focused gene expression to measure quantitative disease activity. Finally, to the best of our knowledge, this is the largest analysis of SLE performed to date that demonstrates that there is a transcriptional signature systemically expressed across different cell types and tissues from patients with SLE and is distinct from other autoimmune and infectious diseases. Our work has the potential to enable more precise molecular definition of SLE that is distinct from other autoimmune diseases.

The role of IFN in SLE has been important in improving the understanding of disease pathogenesis, leading to many publications defining the mechanisms of IFN in SLE(2–5) and several promising clinical trials testing anti-IFN treatments in SLE patients(46, 47). To explore beyond this existing knowledge about the role of IFN in SLE, we specifically separated the SLE MetaSignature into genes related to IFN and genes that were independent of IFN based on a meta-analysis of 16 transcript profiling datasets from IFN-stimulated human cells. We found that the non-IFN SLE MetaSignature was equally accurate in identifying SLE patients. Notably, the non-IFN SLE MetaSignature had higher correlation with SLE disease activity compared to the IFN SLE MetaSignature. Prior studies have likely focused on the IFN-inducible signature due to the high effect sizes of these inflammatory genes. Excluding highly differentially expressed IFN-inducible transcripts allowed us to focus on genes representative of the more nuanced biology underlying SLE.

Neutrophils also play a critical role in SLE pathogenesis. Low density granulocytes serve as the primary source of pro-inflammatory neutrophil extracellular traps (NETs)(3, 36–41). We found that the non-IFN SLE MetaSignature was most elevated in mature neutrophils, which contrasted with the more immature neutropoeisis signature observed in Bennett*, et al*.(3). The non-IFN SLE MetaSignature was also elevated both in low density granulocytes and in response to NETosis-inducing stimulation. Overall, our work further refines the signature of neutropoeisis in SLE, and reinforces an important role for low density granulocytes and NETosis in SLE. Although B cells have an established role in SLE, the SLE MetaScore exhibited mixed results on available sorted B cell gene expression datasets. Due to limited availability of data, we cannot conclusively evaluate whether these challenges are the result of experimental conditions or a lack of signal in B cells.

One of the most exciting discoveries in the SLE MetaSignature is the identification of 14 genes (termed Under-appreciated SLE MetaSignature Genes) that are unrelated to type I IFN or neutrophil-specific gene dysregulation, and are genes that by and large have not previously been implicated in SLE pathogenesis. These newly identified genes fall into categories that include genes with interesting known biology that are expressed in immune cells (e.g., *KLRB1*, *GPR183* (also called *EBI2*), *CD1C*, and *ELANE*); genes involved in inflammation and cellular stress responses (*MT1E*, *MT1F*, and *MT1HL1*); and individual genes related to vitamin B12 metabolism (*TCN2*) and epidermal cellular integrity (*DSC1*).

The most striking group of genes that we identified were members of the metallothionein gene family (*MT1E*, *MT1F*, and *MT1HL1*). Metallothioneins are intracellular, cysteine-rich, metal-binding proteins involved in diverse intracellular functions that include clearance of heavy metals (cadmium, zinc, and copper) from cells, and maintenance of essential ion homeostasis(48, 49). Metallothioneins normally bind zinc(50), an important element and potent antioxidant that influences redox state, enzyme activity, gene transcription, energetic metabolism, cell cycle, cell migration, invasivity, apoptosis, and proliferation(51). Both human cell line and animal studies have indicated a role for metallothioneins in protection against cadmium toxicity(52–55). Metallothioneins can be activated by a variety of stimuli, including metal ions, cytokines and growth factors, as well as oxidative stress and radiation(51, 56). During oxidative stress, metallothioneins are up-regulated to protect the cells against cytotoxicity, radiation, and DNA damage(57–59). Interestingly, metallothionein proteins are expressed at elevated levels in the kidneys of lupus nephritis patients(60). We found that transcript profiles of human cell lines exposed acutely or chronically to heavy metals resembled the Under-appreciated SLE MetaSignature. We therefore hypothesize that up-regulation of metallothioneins in SLE may be a protective response to elevated oxidative stress during chronic inflammatory responses, and/or exposure to environmental sources of heavy metals found in tobacco, particulate air pollutants (e.g., traffic-derived particles from tires and automative brakes), diet, and industrial waste(61–64). The importance of metallothioneins in SLE pathology is underscored by the observation that two additional family members (*MT1A* and *MT2A*) are induced by interferons and were identified in the 93 gene SLE MetaSignature.

The *ELANE* gene encodes neutrophil elastase (NE), a serine protease implicated in host defense and tissue injury. In addition to elastin, NE also hydrolyzes proteins within azurophil granules, extracellular matrix proteins, the outer membrane protein A (*OmpA*) of *E. coli*, and the virulence factors of other bacteria(65). In contrast to the digestive serine proteases, NE has unusually high affinity for nucleic acids(66). In naive neutrophils, NE is normally stored in azurophilic granules(67, 68). Upon activation, NE translocates from azurophilic granules to the nucleus, where it partially degrades specific histones, thereby promoting chromatin decondensation and regulating the formation of NETs(69). NE knockout mice are susceptible to bacterial and fungal infections(70, 71). Mutations in *ELANE* can lead to cyclic and severe congenital neutropenia(72). Furthermore, the NE enzyme may also play a role in various lung, bowel, and skin inflammatory diseases(73). Dysregulation of *ELANE* in SLE was previously noted in a single cohort gene expression profile(74). Although known as a neutrophil-expressed gene (hence its name), we did not identify neutrophil-specific dysregulation of *ELANE* in our analysis of SLE datasets, and rather than being classified under the neutrophil-related SLE MetaSignature genes, *ELANE* was instead classified as a “Under-appreciated SLE MetaSignature” gene. Unexpectedly, our further analysis of cell type expression of *ELANE* using immunoStates(35) indicated that *ELANE* is most differentially expressed in hematopoietic progenitor cells and basophils [Figure S8]. This suggests that novel functions for *ELANE* in other cells, in addition to neutrophils, may be involved in the pathophysiology of SLE.

*DSC1* encodes a calcium-dependent glycoprotein in the desmocollin subgroup of the cadherin family. The desmocollins are critical adhesive proteins of the desmosome cell-cell junction linking epithelial cells, and are required for cell adhesion and desmosome formation. *DSC1* is expressed in the upper epidermis of the skin(75) and has been implicated as an autoantigen for bullous skin disease(76, 77), which is also frequently manifested in SLE patients(77). Mice lacking *DSC1* exhibit epidermal fragility accompanied by defects in epidermal barrier and differentiation(78). Neonatal mice lacking desmocollin develop epidermal lesions, and older mice develop ulcerating lesions resembling chronic dermatitis. Based on the above observations, we speculate that the abnormally low levels of *DSC1* in SLE patients lead to reduced adhesion and barrier maintenance of the upper epidermis, increasing the susceptibility to development of bullae and dermatitis.

*KLRB1* (also known as *CD161*) encodes a C-type lectin-like receptor that is composed of a disulfide-linked homodimer of ∼40 kDa subunits, and is part of the natural killer gene complex (NKC)(79). *KLRB1* has been previously shown to be down-regulated in SLE(37, 80, 81). This gene is expressed by NK cells, subsets of αß and γδ T cells, and invariant CD1d-specific Natural Killer T (NK-T) cells(82–84). The *KLRB1* receptor, by interacting with its ligand *LLT1*(85, 86), plays an inhibitory role in NK cell-mediated cytotoxicity and IFNγ secretion during immune responses to pathogens(82, 85–87). Polymorphisms in *KLRB1* are associated with structural alterations of the protein, and impact its regulatory functions on NK cell homeostasis and activation(88). In contrast to its inhibitory potential in NK cells, the function of *KLRB1* in T cells is less clear, with reports suggesting both co-activating(83, 87, 89) and inhibitory(90, 91) effects. *CD161* (*KLRB1*) has been used as a marker to define Th17 and Tc17 subsets of CD4+ and CD8+ T cells that secrete the proinflammatory cytokine interleukin IL-17. However, a more recent study found that *CD161*-expressing T cell subsets are not all committed to the Th17 axis but are much more diverse, and that expression of *CD161* identifies a transcriptional and functional phenotype shared across human T lymphocytes that is independent of both T cell receptor (TCR) expression and cell lineage(89). The dysregulation of *KLRB1* in SLE may be directly linked to aberrant IFNγ signaling pathways and immune cell subpopulations in this disease.

*GPR183* (also known as *EBI2*) encodes the G Protein-Coupled Receptor 183 that binds oxysterols, the most potent of which is 7α, 25-dihydroxycholesterol (7α,25-OHC)(92). GPR183 is up-regulated in a Burkitt’s lymphoma cell line upon Epstein-Barr virus infection(93), an infection that is also strongly linked to SLE(93, 94). Interestingly, *GPR183* is also strongly induced in UVB-irradiated skin biopsies(95) and UV light has been postulated to induce SLE photosensitivity(96) and DNA damage-driven apoptosis(97). The *GPR183* protein is a negative regulator of IFN(98). In lymphoid organs, *GPR183* plays a key role in mediating the migration and antibody response of multiple immune cell types including B cells, T cells, dendritic cells and monocytes(99–103). *GPR183*-deficient mice have fewer plasma cells, reduced antibody titers(99, 100), and diminished CD4+ splenic dendritic cells. In another study, mice lacking *GPR183* or its 7α,25-OHC ligand show defects in the trafficking of group 3 innate lymphoid cells (immune cell subsets enriched in the intestine) and defects in lymphoid tissue formation in the colon(104). *GPR183* has been implicated in inflammatory and autoimmune diseases, including multiple sclerosis(105), inflammatory bowel disease(106), Crohn’s disease(106), type 1 diabetes, and cancer(103). In multiple sclerosis, data from the experimental autoimmune encephalomyelitis (EAE) animal model suggest that *GPR183* is a critical mediator of CNS autoimmunity and regulates the migration of autoreactive T cells into inflamed organs(107). Thus, the intriguing links between *GPR183* and SLE through Epstein-Barr virus, IFN, and UV-light, as well as its important functions in instructing immune cell localization and antibody response, identify *GPR183* and its ligand 7α, 25-OHC as potential biomarkers and/or therapeutic targets for SLE.

Research in many immunological disorders, including SLE, has recently focused on the importance of immunometabolism (immune related dysregulation of metabolic pathways) in disease(108, 109). In SLE, particular focus has been on T cell metabolism (mitochondria, oxidative stress, mTOR, glucose, and cholesterol pathways), with additional interest in B cells (glycolysis and pyruvate), macrophages (stress response), dendritic cells (mTOR, fatty acids), and neutrophils (NETosis, oxidation)(109). Concordant with these prior findings, our pathway analysis(28) of the SLE MetaSignature recapitulated many similar immunometabolic pathways in SLE, including pyruvate metabolism, fructose galactose metabolism, and oxidative stress response. In addition, our pathway analysis identified many non-inflammatory pathways involved in nucleic acid metabolism (including formyltetrahydrofolate biosynthesis, salvage pyrimidine deoxyribonucleotides, salvage pyrimidine ribonucleotides, and purine metabolism). Two therapies in SLE, methotrexate(110) and leflunomide(111–113), both inhibit nucleic acid metabolism(114–117), and other new molecular entities that target these pathways are entering clinical trials. Collectively, our pathway analysis results reinforce the importance of immunometabolic pathways in SLE pathogenesis.

Arguably, our approach, which leverages heterogeneity within patient populations to identify a common transcriptional signature across all patients with SLE, is ill-suited in the era of “personalized medicine”. A goal of personalized medicine is to cluster heterogeneous patients into homogeneous subgroups, which does not account for the individual variations that should be targeted. The underlying assumption in this approach is that the individual variation between subgroups is likely causal, which can be targeted to improve therapy and outcomes. However, it is equally likely that the disease-causing biology may be the same across all patients, and the variation observed between patients and subgroups is a result of environmental exposures (e.g., epigenetics). For example, studying a homogeneous patient population (i.e., a subgroup of patients) may identify a signature that explains the variation between groups but may not be causal. In this case targeting the variation between patients would not be sufficient to effectively treat the disease. Therefore, we believe that a more suitable approach would be to complement “personalized medicine” with “precision medicine” in SLE such that it first provides a precise molecular definition of SLE, such as we have done here. This could then lead to identification of multiple drug targets and corresponding therapies, increasing the number of drugs available to treat SLE patients. Only 1 drug has received FDA-approval in the last 60 years for treatment of SLE. Following the precise molecular definition of SLE, “personalized medicine” could help to identify which patients will respond to which of these drugs, given they have molecularly defined SLE. Previously, we have shown utility of such an approach across a number of disease domains that conserved signatures of disease can be vital to identifying novel mechanisms, and treatments of a disease(13, 16, 24, 118).

Between our own efforts and those of other researchers, we hope that the SLE MetaSignature is used to guide future experiments and analyses to illuminate the functional implications of each gene/protein. We anticipate that the full SLE Metasignature, and particularly the 14 gene subset that is interferon independent and unrelated to neutrophils, will be tested in blood and tissue derived from prospectively collected SLE cohorts to determine if a relationship can be identified between SLE flares, clinical subgroups, and responses to newly-tested therapies. Another important question to be addressed is whether the proteins encoded by these genes are similarly expressed at abnormal levels or in unanticipated cell populations, and if so, in which tissues. Our results will help guide targeted analyses of ongoing studies in SLE blood and kidney samples using single cell technologies such as scRNA-Seq, ATAC-Seq, Cytometry Time of Flight (CyTOF), Multiplexed Ion Beam Imaging (MIBI), and CO-Detection by IndEXing(119). Many of these novel methods are being utilized by the Accelerating Medicines Partnership RA/SLE program to characterize human SLE tissue, with a goal to identify novel pathways and disease targets(120). Ongoing studies in our group using CRISPR screens and immunohistochemistry are interrogating the role played by each of these genes in cultured immune cells, as well as the effect of the 14 gene subset on interferon signaling, neutrophil biology, and animal models.

Our analysis has a few limitations. First, we focused on identifying a gene signature that is conserved between cohorts and across samples, and does not identify patient subgroups. While this is beneficial for capturing features that are consistent across populations, it is ill-suited for identifying subgroups of disease. Second, because we only used publicly available datasets, our analyses were restricted to the comparisons available in the public data, including tissues, cell types, and diseases sampled. To enable even richer analysis, we encourage the research community to contribute richly annotated datasets to the public domain. In the context of SLE, particularly important annotations, when available, include: age, sex, SLEDAI with individual components specifically recorded, drugs at the time of blood draw, drug doses and start dates, organ system involvement, cell proportions from complete blood count or flow cytometry.

Recent studies have been dominated by important discoveries that link type I IFN, neutrophils, and NETs to SLE. We have identified a unified SLE MetaSignature that implicates 14 under-appreciated genes in SLE pathogenesis, only four of which were identified through a direct PubMed search in the context of SLE (KLRB1, GRN, CD1C, and ABCB1, with 2, 5, 7, and 9 references each, respectively). More thorough scouring of published literature reveals connections to additional genes, including ELANE, EBI2, and LHFPL2, but none of these genes have garnered significant attention in the context of SLE. Included in this set of 14 genes are eight genes described above that have plausible roles in SLE, because they are expressed in immune cells or skin, or are associated with stress responses. Perhaps even more interesting are the six genes (*ABCB1*, *GRN*, *LHFPL2*, *NAP1L3*, *TCN2*, and *VSIG1*) that are not linked to the immune system or plausible pathogenic mechanisms, and have few if any associations with known autoimmune diseases. It is well known that scientists fall prey to the “streetlight effect” - looking for answers where the light is better rather than where the truth is more likely to lie(121–124). While many of the genes we have identified in the Under-appreciated SLE MetaSignature “make mechanistic sense” based on the literature, and will be candidates for immediate characterization because of what we already know about them, we should not lose sight of the six new genes which had previously been in the shadows but are now illuminated.

## Methods

### Gene expression meta-analysis

We used R package MetaIntegrator for integrating discovery cohorts as described previously(125) to identify differentially expressed genes between healthy controls and patients with SLE. First, we computed an effect size for each gene in each study as Hedges’ adjusted g. Next, we summarized the effect sizes across all studies for each gene using DerSimonian-Laird method for random effects inverse variance model, where each effect size is weighted by the inverse of the variance in that study. Finally, we corrected the p-values for the summary effect size of each gene for multiple hypothesis testing using Benjamini-Hochberg false discovery rate (FDR)(126).

Briefly, we downloaded gene expression microarray data from 40 independent experiments with 7,471 samples from the Gene Expression Omnibus (GEO)(127). Accession numbers, tissue of origin, data center, sample size, and PubMed identifiers (IDs) for all datasets are listed in Table 1. We randomly selected 6 datasets composed of 370 whole blood and PBMC samples as “Discovery” datasets, based on our previous finding that 4-5 datasets with 250-300 samples have sufficient statistical power to find a robust reproducible disease gene signature using our multi-cohort analysis framework. The six experiments used in the discovery set were required to have both SLE patients and healthy volunteers for use as cases and controls, respectively, and were limited to analyses of PBMCs or blood. Based on previous analyses, a combined sample size of 370 patients from six datasets was considered sufficient for the discovery phase of the meta-analysis(26). We used eight gene expression datasets with 2,407 samples from individuals who have SLE and healthy volunteers in PBMCs or whole blood as a validation set. The remaining 26 datasets and 4,694 samples, which include sorted cell and tissue data, were assigned to the extended validation set.

We used the following thresholds in our meta-analysis of the discovery set to select genes in the SLE MetaSignature: absolute value of effect size greater than one; false discovery rate less than five percent; and measurements of individual genes in the identified signature in at least four datasets. We then used the genes in the SLE MetaSignature to calculate an SLE MetaScore for each patient sample. To avoid an overrepresentation of up or down-regulated genes in the SLE MetaScore, we modified the default MetaIntegrator calculateScore function. The modified function calculated the geometric mean of the up-regulated genes and the inverse of the down-regulated genes (rather than subtracting the geometric mean of the down-regulated genes from the geometric mean of the up-regulated genes). We then used the z-score to scale the geometric means across samples in the same study.

### SLEDAI correlation significance

We generated 100 random gene sets with the same number of positive and negative genes as the SLE MetaSignature. We evaluated the correlation for each of these gene sets with SLEDAI using the same method as the SLE MetaScore. For each randomization, we evaluated the median correlation across all 5 studies with SLEDAI measurements.

### IFN meta-analysis

We downloaded gene expression microarray data from 15 datasets [Table S2] of primary human cells stimulated with Type I IFN from GEO(25) using the MetaIntegrator R package(15). Using MetaIntegrator, we ran meta-analysis with unstimulated cells treated as controls and type I IFN stimulations as cases. In our analysis, we utilized the effect size and effect size FDR estimates for IFN based on our meta-analysis.

### Neutrophil, NK cell, and heavy metal datasets

We identified 7 relevant datasets for our follow-on analyses of the SLE MetaSignature and downloaded them from GEO(25) using the MetaIntegrator R package(15). These datasets included stimulations to induce NETosis, intermediate cell populations from neutropoeisis, and heavy metal exposures in cell lines.

### Pathway analysis

Pathway analysis was performed using the Differential Expression Analysis for Pathways (DEAP) tool(28). As input, gene effect size measurements were used from all discovery and validation datasets. 1000 random rotations of the data were performed to assess statistical significance. Pathways were downloaded from the PANTHER pathway database(128).

### Stanford pSLE patient cohort

All subjects were recruited and all samples were collected following protocols approved by the Stanford University Institutional Review Board (IRB protocol #13952, 14734). Patients who fulfilled American College of Rheumatology (ACR) revised diagnostic criteria for SLE were consented at the Pediatric Rheumatology Clinic at Stanford Children’s Health Lucile Packard Children’s Hospital (LPCH), Stanford(129). Age-appropriate consent and assent was obtained. A total of 43 new-onset SLE patients were recruited along with 12 patients diagnosed with Juvenile Idiopathic Arthritis (JIA) as disease controls. Initial whole blood samples were obtained within a mean of 5 days of diagnosis. One patient (93) initially only had 3 ACR criteria, but was monitored and diagnosed with SLE (meeting 4 ACR criteria) ∼3 years after the initial sample was obtained. This patient was not included in the comparison of SLE versus JIA. Clinical assessment of disease activity and treatment was conducted using a modified Safety of Estrogens in Lupus Erythematosus National Assessment Systemic Lupus Erythematosus Disease Activity Index (SELENA-SLEDAI(130)), calculated for each visit. Whole blood samples were collected from 10 healthy volunteers.

### Stanford pSLE cohort sample collection and RNA processing

At each patient visit, approximately 3ml of whole blood was collected into a Tempus™ Blood RNA Tube (Life Technologies #4342792) and frozen at −20°C for a minimum of 30 days. Batched samples were thawed and processed for RNA extraction using a Tempus™ RNA Isolation Kit (Life Technologies #4380204). All RNA samples were stored at −80°C prior to use. Samples were analyzed using a Bioanalyzer (Agilent Technologies) to confirm RNA quality prior to assay. All samples had RNA Integrity Numbers (RIN) > 7.9.

### Fluidigm Transcript Analysis

A panel of 33 genes was selected from the genes in the SLE MetaSignature, based on availability of TaqMan probes and meta-analysis effect size in a preliminary version of the SLE MetaSignature. Once we finalized the SLE MetaSignature, we used those genes which were in the final SLE MetaSignature, and for which we had acquired probes based on the preliminary analysis. TaqMan probes are listed in Table S4.

The Human Immune Monitoring Center (HIMC) at Stanford University performed transcript analysis. Reverse transcription was conducted at 50°C for 15 min using the High Capacity Reverse Transcription kit (ABI) and 10-50 ng total RNA. Pre-amplification of cDNA was performed using the TaqMan PreAmp Master Mix Kit (Invitrogen). TaqMan probes are listed in Table S1. Reverse transcriptase was inactivated and Taq polymerase reaction was initiated by bringing samples to 95°C for 2 min. The cDNA was preamplified by denaturing for 10 cycles at 95°C for 15 s, then annealing at 60°C for 4 min. The cDNA product was diluted 1:2 with 1x TE buffer (Invitrogen). 2X Applied Biosystems Taqman Master Mix, Fluidigm Sample Loading Reagent, and preamplified cDNA were mixed and loaded into the 48.48 Dynamic Array (Fluidigm) sample inlets, followed by 10X assays. Real-time PCR was carried out with the following conditions: 10 min at 95°C, followed by 50 cycles of 15 s at 95°C and 1 min at 60°C. All reactions were performed in duplicate. Gene Ct values were normalized to GAPDH, Beta-actin, and Beta-2-microglobulin. Average DCt values of controls were then subtracted from target gene DCt values to give ddCT. Relative gene expression levels were calculated as 2^-ddCt^.

### Statistical power analysis

We computed statistical power for the meta-analysis of gene expression between healthy controls and patients with SLE in both discovery and validation analysis(131) under no, low, moderate, or high heterogeneity assumption. In discovery datasets, we had >90% statistical power at p-value of 0.01 to detect effect size >0.44 and >0.9 when assuming no heterogeneity or high heterogeneity, respectively. In validation cohorts, we had >90% statistical power at p-value of 0.01 to detect effect size >0.3 and >0.59, when assuming no or high heterogeneity, respectively. In the cohort of whole blood samples from pediatric patients with SLE used for validation using Fluidigm, we had statistical power >99%.

### Code and data availability

All code and data necessary for reproducing this analysis are available at https://wiki.khatrilab.stanford.edu/sle.

### Statistics

The statistics related to gene expression meta-analysis, correlation analysis, pathway analysis, and statistical power analysis are all described in the appropriately title sections above. In general, a Benjamini-Hochberg False Discovery Rate (FDR) threshold of 0.05 was used to label any findings as “significant” in this manuscript.

### Study approval

The vast majority of the data was obtained from public repositories. For the prospective validation analysis, all subjects were recruited and all samples were collected following protocols approved by the Stanford University Institutional Review Board (IRB protocol #13952, 14734).

## Author contributions

Designing experiments: WAH, PJU, PK; Conducting experiments: DJH, VKD, IB; Acquiring data: WAH, EB, CRB; Analyzing data: WAH, DJH, EB, CRB; Acquiring samples: IB, PJU; Writing manuscript: WAH, DJH, VKD, EB, GY, IB, CRB, RM, PJU, PK

## Acknowledgements

We would like to thank the pediatric SLE and JIA patients and their families; the healthy volunteers who participated in this study; HIMC for performing Q-PCR assays; and members of the Khatri and Utz labs.

WAH is funded by the National Science Foundation Graduate Research Fellowship under Grant No. DGE-114747 and NIH NLM T15 LM 007033. IB was supported by the National Institutes of Health (NIH) (grant number K08-AI-080945), the Stanford Child Health Research Institute (Child Health Research Program Pilot Grant for Early Career Investigators), and the Arthritis Foundation (Postdoctoral Fellowship). E.B. was supported by the Stanford Gabilan Graduate Fellowship in Science and Engineering and the Stanford Women and Sex Differences in Medicine (WSDM) Seed Grant. GY was supported by the Stanford Medical Scientist Training Program (NIGMS grant# 5T32GM007365-43). PK is funded by the Bill and Melinda Gates Foundation, and the National Institute of Allergy and Infectious Diseases grants 1U19AI109662, U19AI057229, U54I117925, and U01AI089859. PJU is the recipient of gifts from the Henry Gustav Floren Family Trust, Elizabeth Adler, and the Baxter Foundation. PJU is supported by NIH grants NIAID U19-AI1110491, Stanford Autoimmunity Center of Excellence (ACE); NIAID 1 UM1AI110498-01, ACE Collaborative Project; 1 U19-AI090019; and NIAID 1 R01 AI125197-01.

---

**Figure S1.**
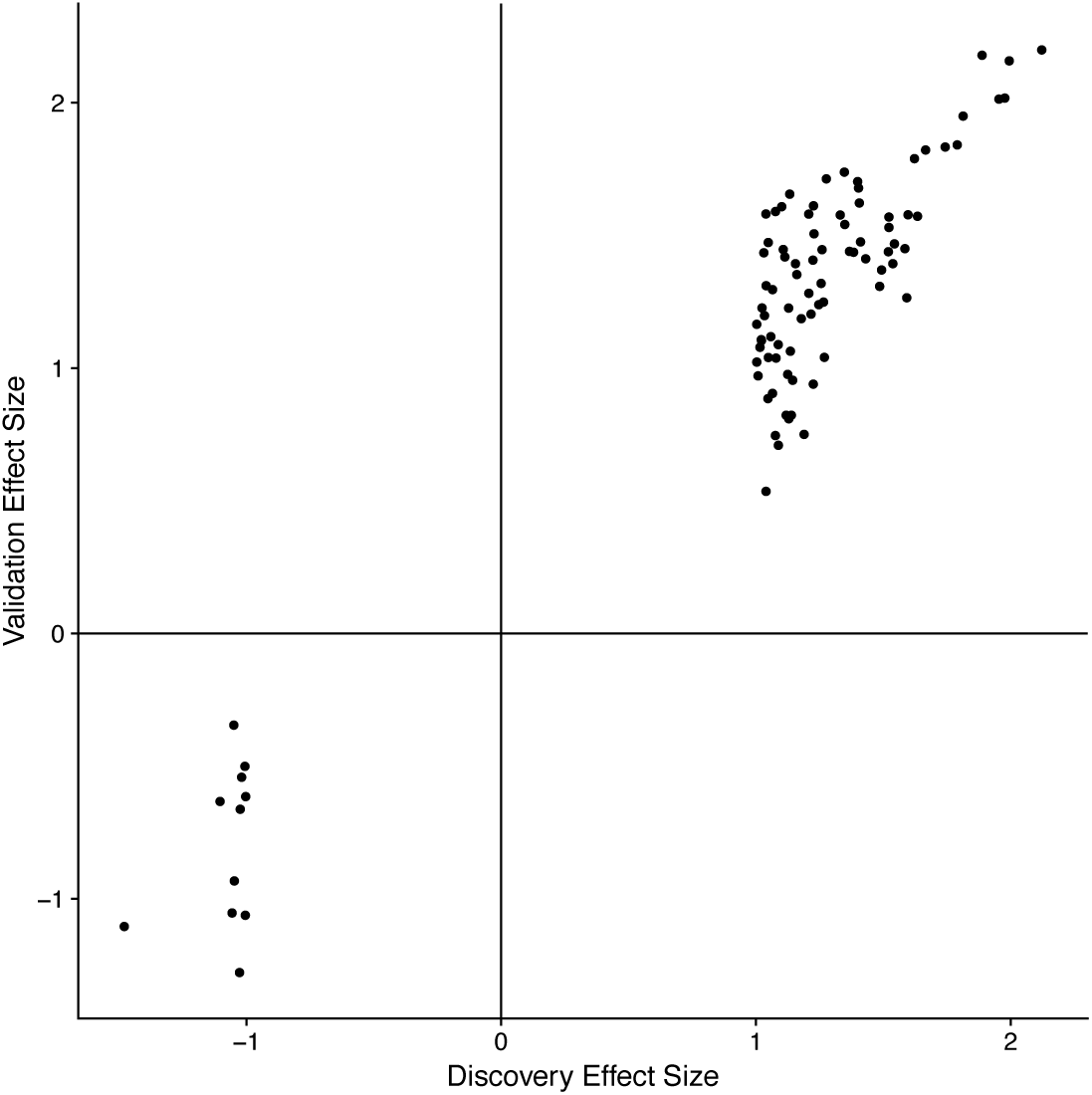
Discovery vs. validation effect sizes for the SLE MetaSignature. Effect sizes for all 93 genes from the SLE MetaSignature are displayed in both the discovery and validation data.

**Figure S2.**
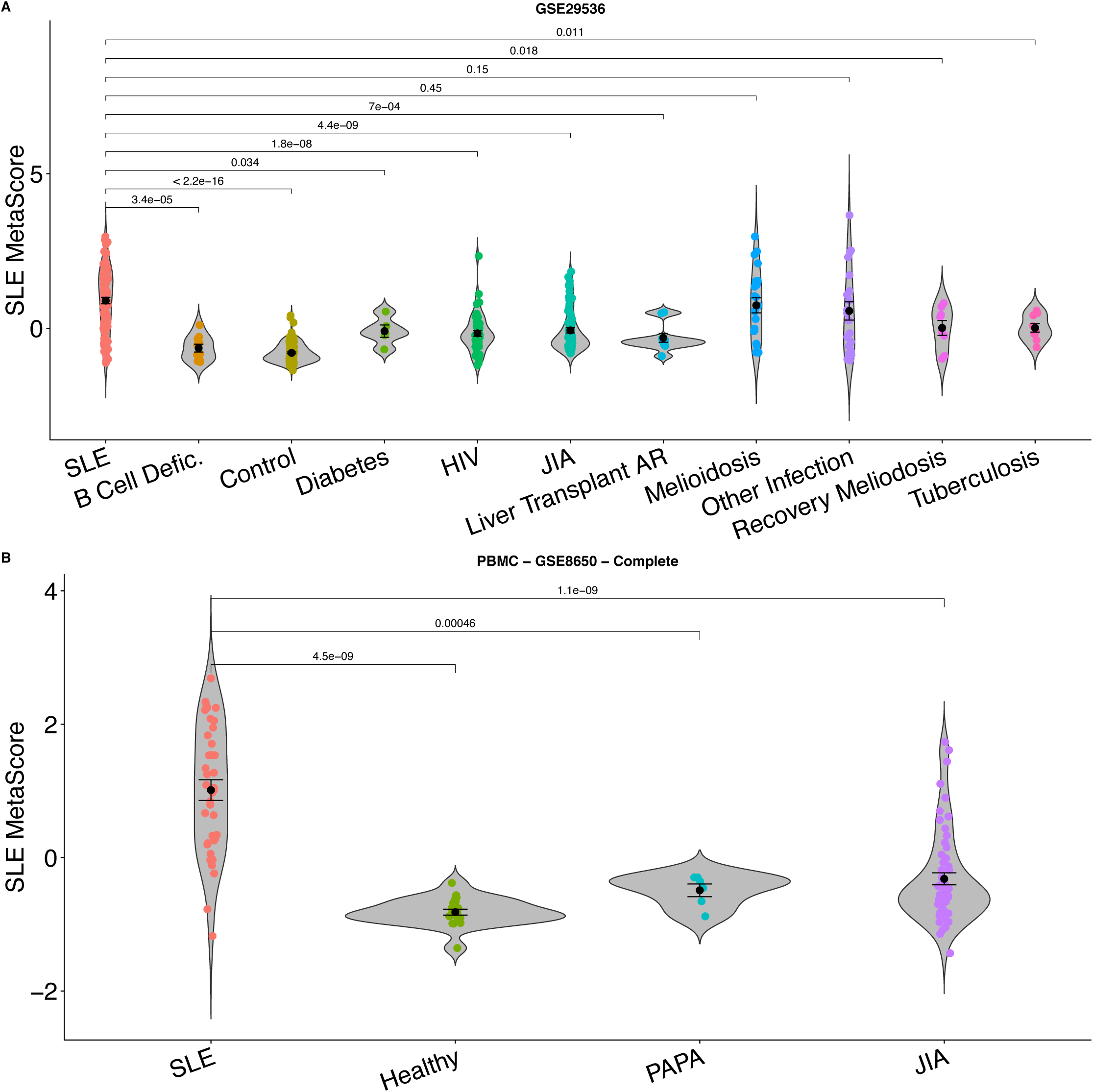
SLE compared to other diseases. Related to Figure 3. (A and B) Violin plots for additional datasets comparing SLE to non-SLE diseases. For all panels: B cell deficiency (B Cell Defic.); type 2 diabetes (Diabetes); human immunodeficiency virus (HIV); juvenile idiopathic arthritis (JIA); liver transplant acute rejection (Liver Transplant AR); pyogenic arthritis, pyoderma gangrenosum and acne (PAPA).

**Figure S3.**
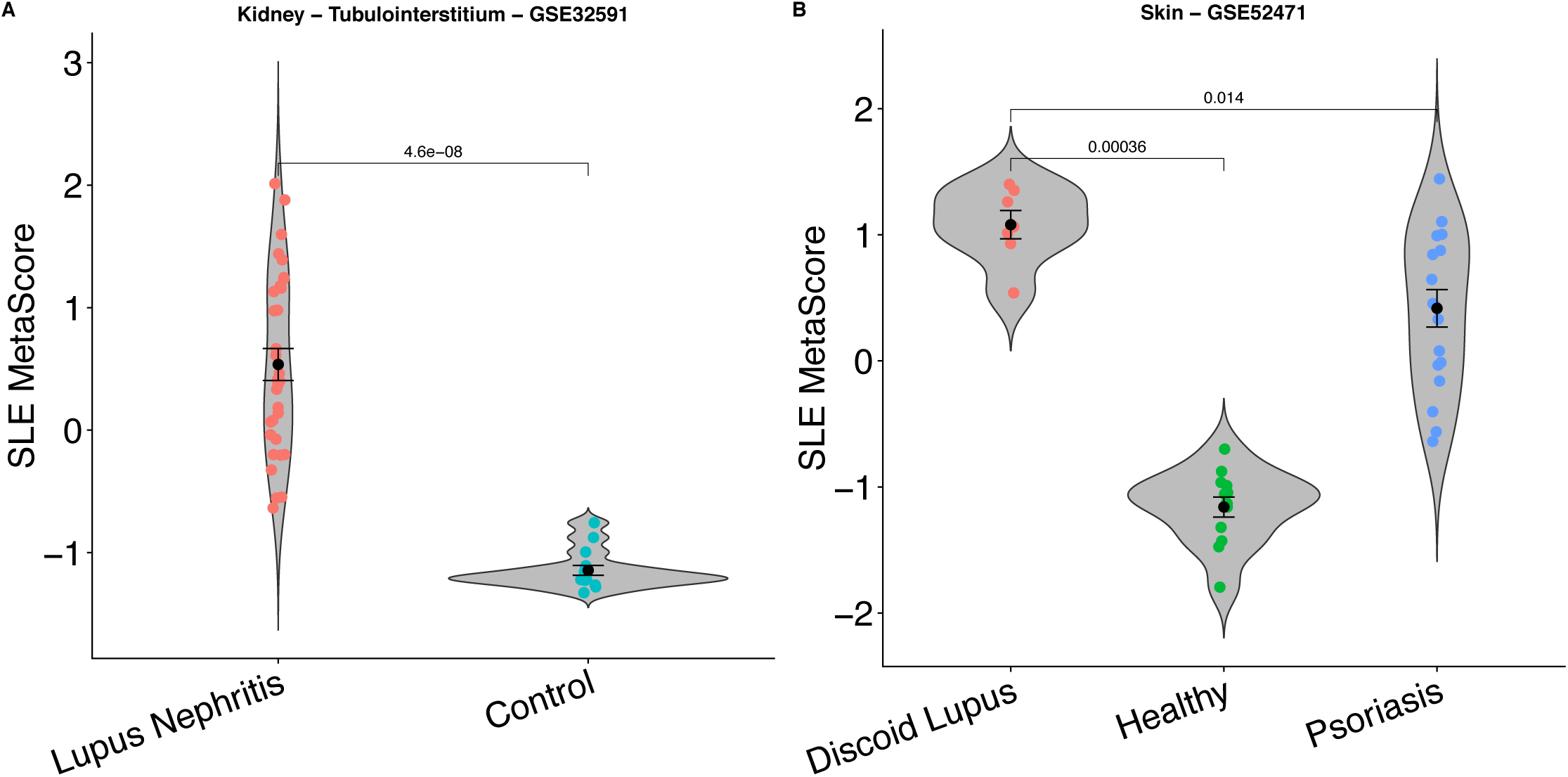
SLE MetaSignature in other tissues. Related to Figure 3. Violin plots for additional datasets examining SLE in kidney tubulointerstitium (A) and skin (B) tissues.

**Figure S4.**
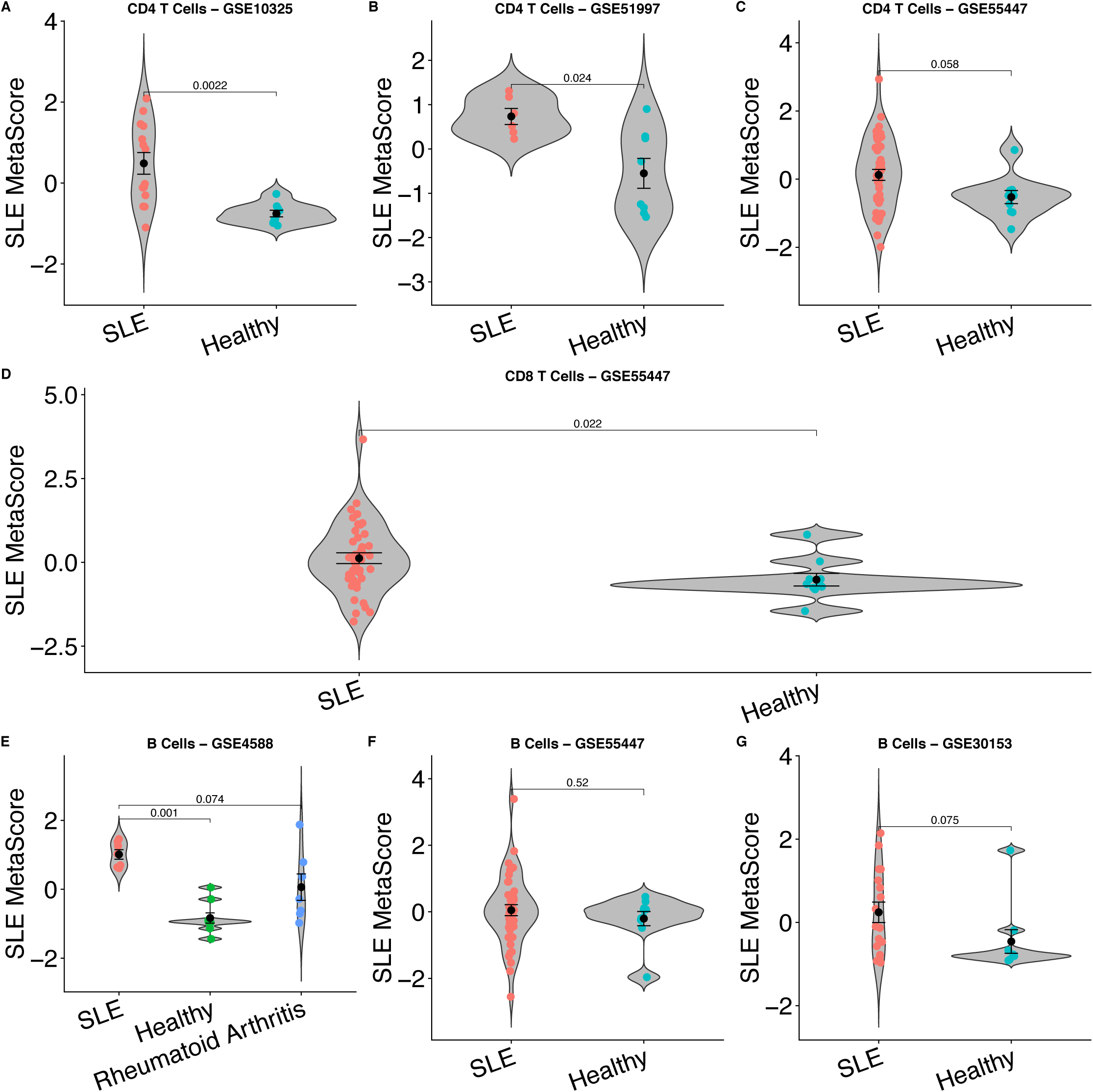
SLE MetaSignature in other cell types. Related to Figure 3. Violin plots for additional datasets examining the SLE MetaSignature in sorted cell populations, including CD4 T cells from different datasets (A-C), CD8 T cells (D), and B cells in different datasets (E-G).

**Figure S5.**
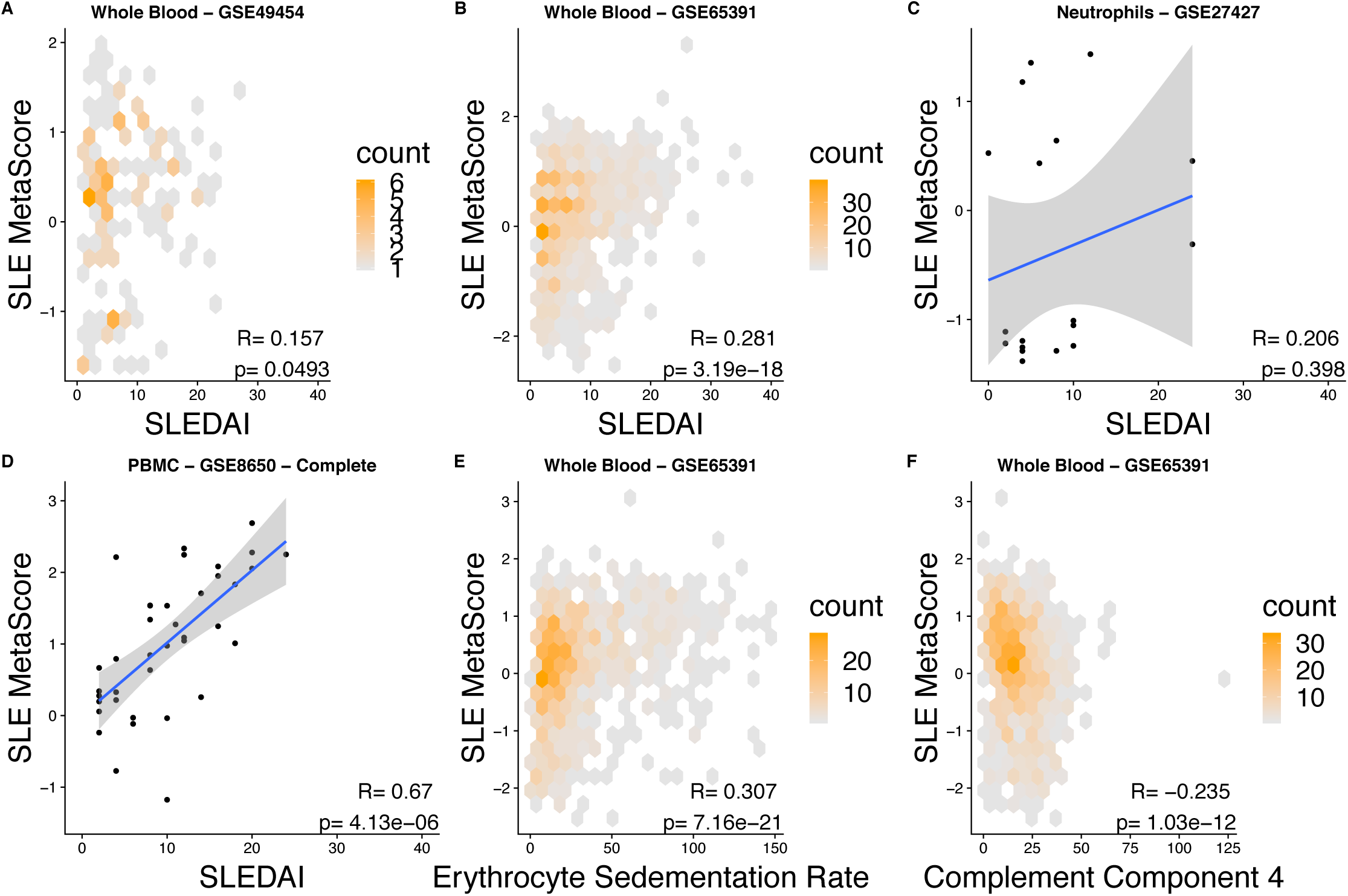
SLE disease activity. Related to Figure 4. (A-D) SLE MetaScore vs. SLEDAI in whole blood (A-B), neutrophil (C), and PBMC (D) samples from SLE patients. (E) SLE MetaScore vs. erythrocyte sedimentation rate. (F) SLE MetaScore vs. complement C4 levels in SLE patients.

**Figure S6.**
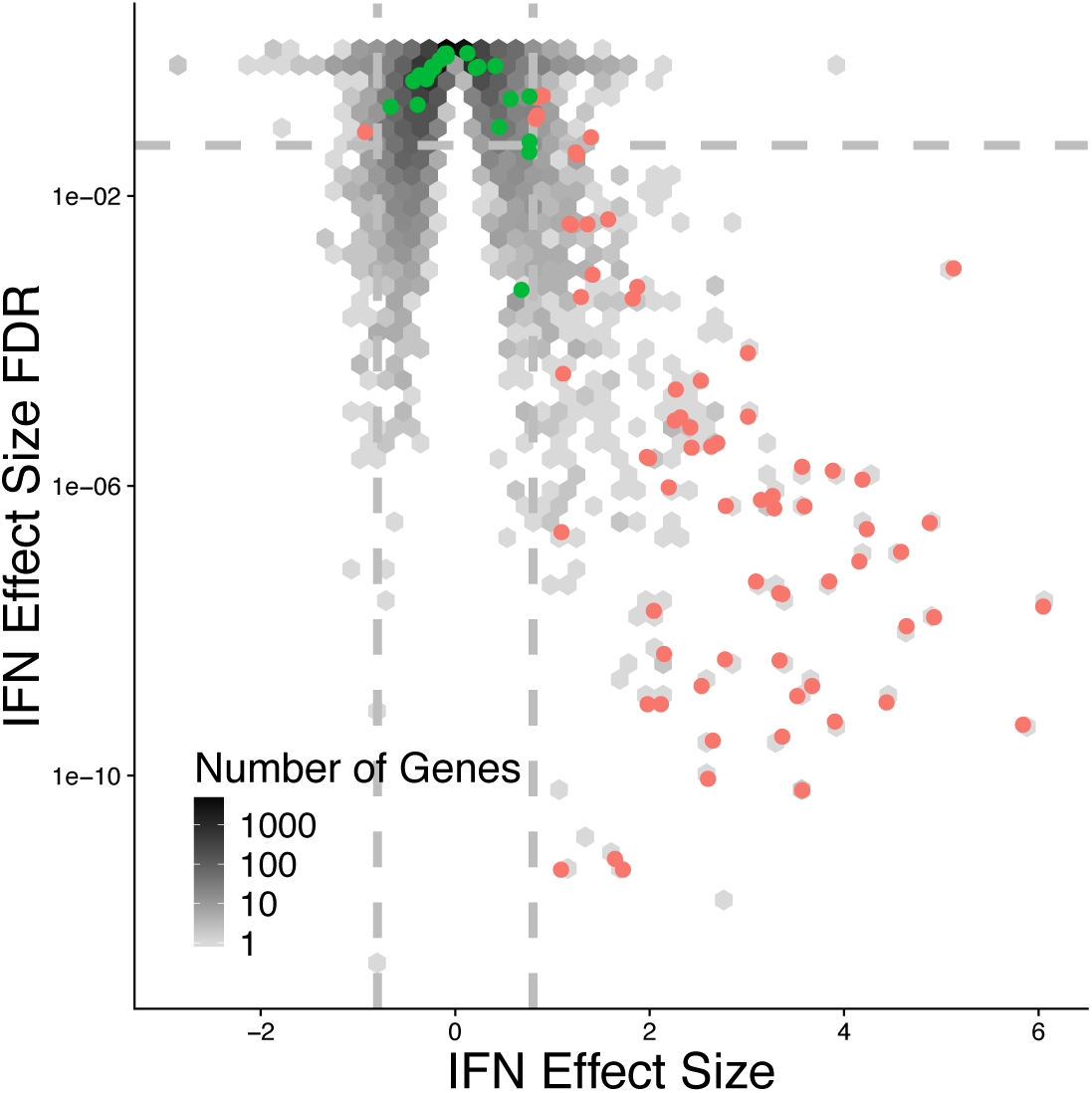
IFN volcano plot. IFN effect size vs. IFN effect size false discovery rate. Red points indicate the 70 genes which are in the SLE MetaSignature and are significantly different in response to IFN. Green points indicate the 23 genes which are in the SLE MetaSignature and are not significantly different in response to IFN.

**Figure S7.**
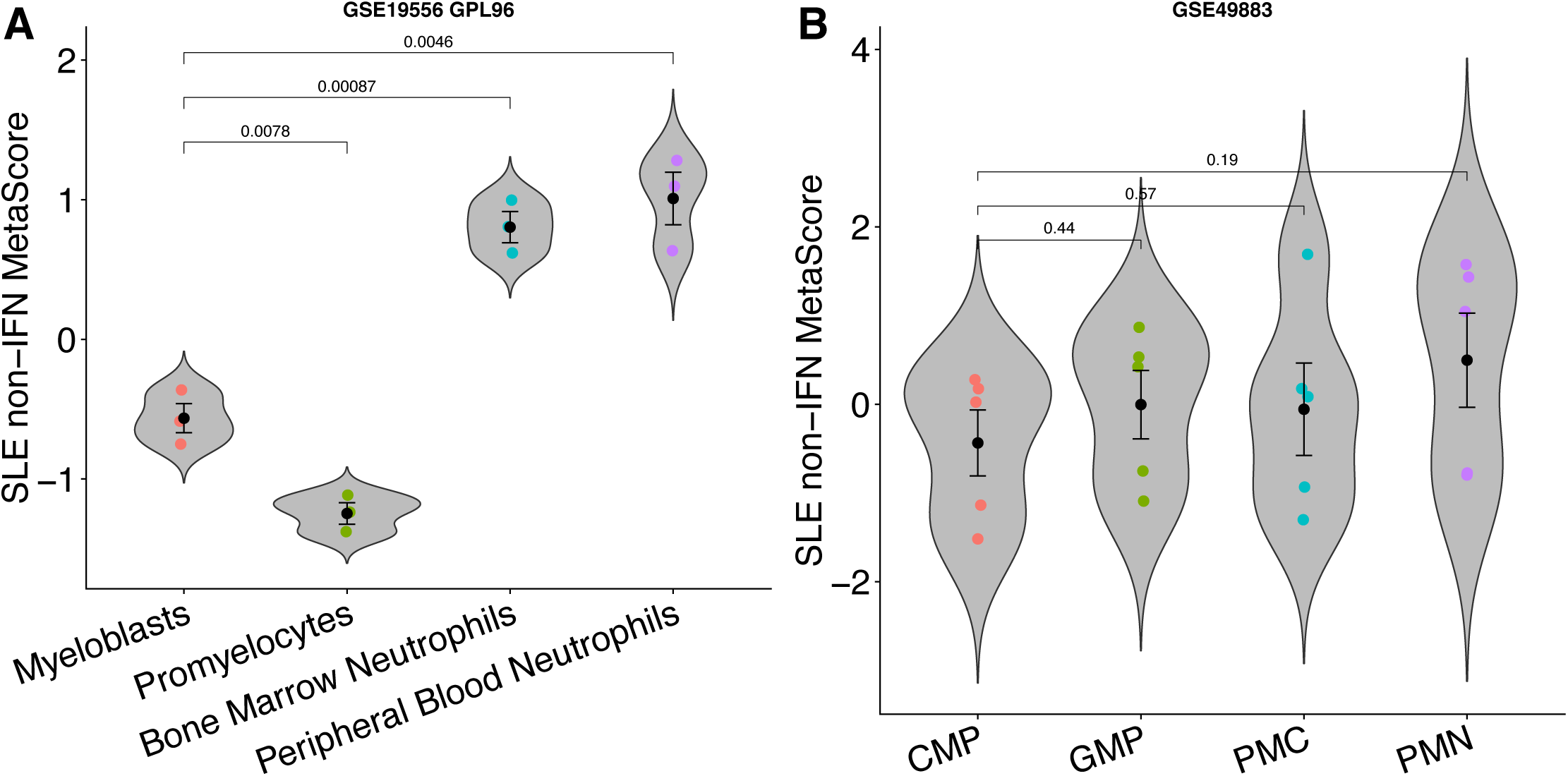
Neutrophil development. Related to Figure 7. SLE non-IFN MetaScore is elevated in mature neutrophils in different datasets of neutropoeisis (A, B).

**Figure S8.**
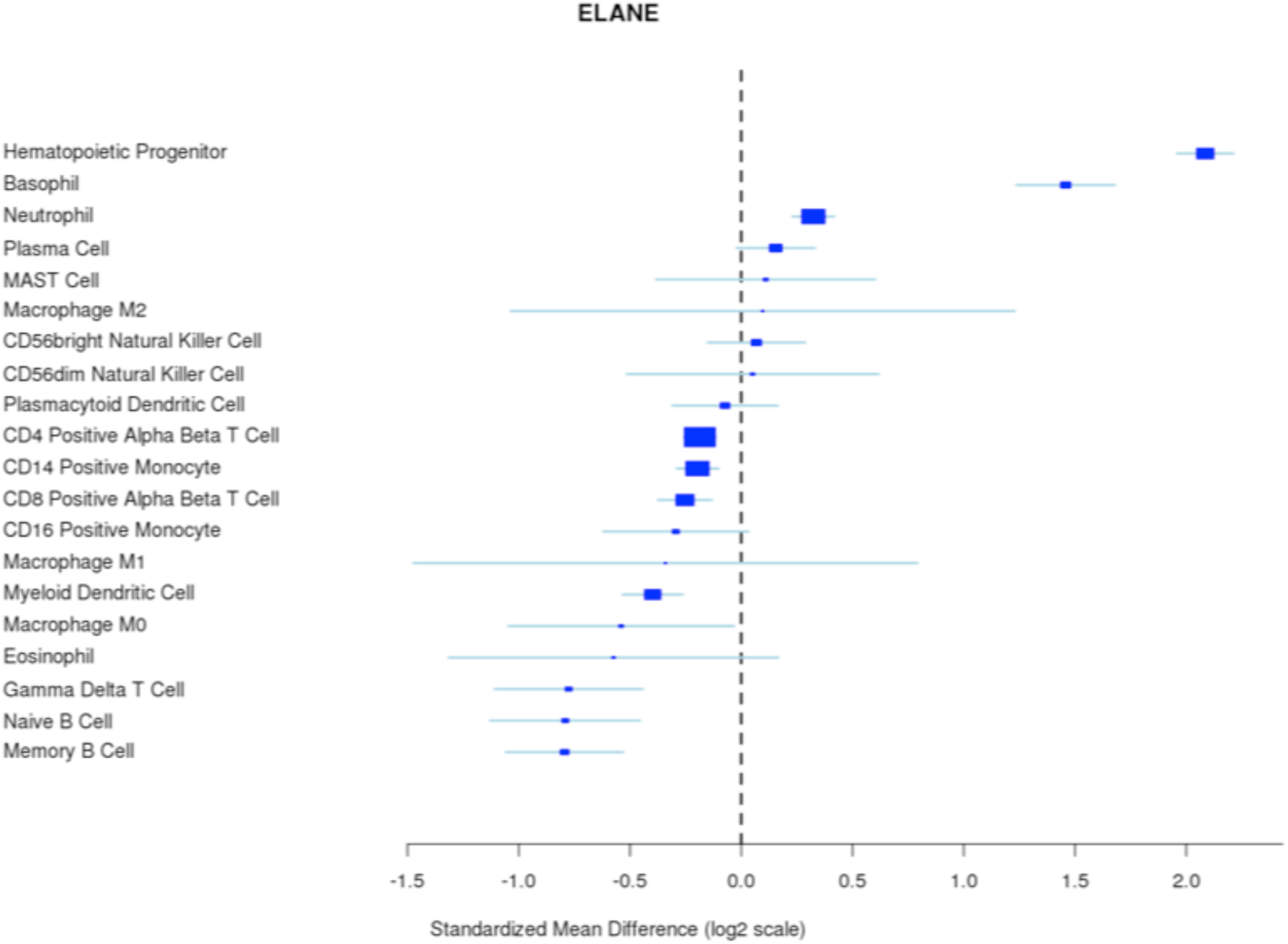
*ELANE* gene effect sizes across cell types. *ELANE* is most differentially expressed in hematopoietic progenitor cells compared to all other cell populations.

**Figure S9.**
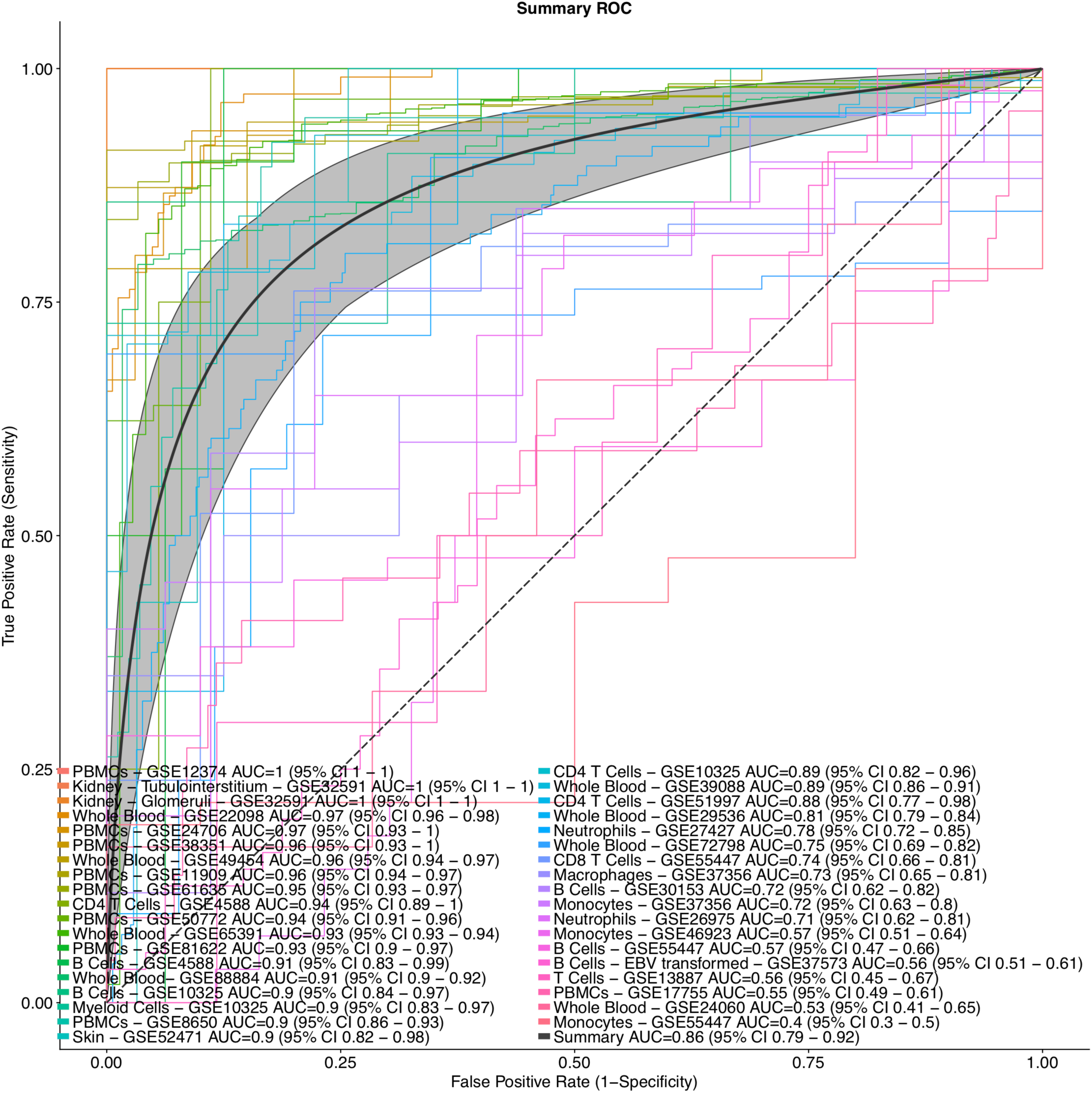
Extended validation ROC. ROC curves for data in the extended validation set. Some datasets also appear in the discovery and validation ROC plots, but may have different AUC estimates, because we have included additional patients which were excluded from the discovery and validation plots due to failure to match our discovery/validation criteria (in which subjects with other diseases were included/excluded, and follow up longitudinal visits from the same subject were included/excluded).

**Figure S10.**
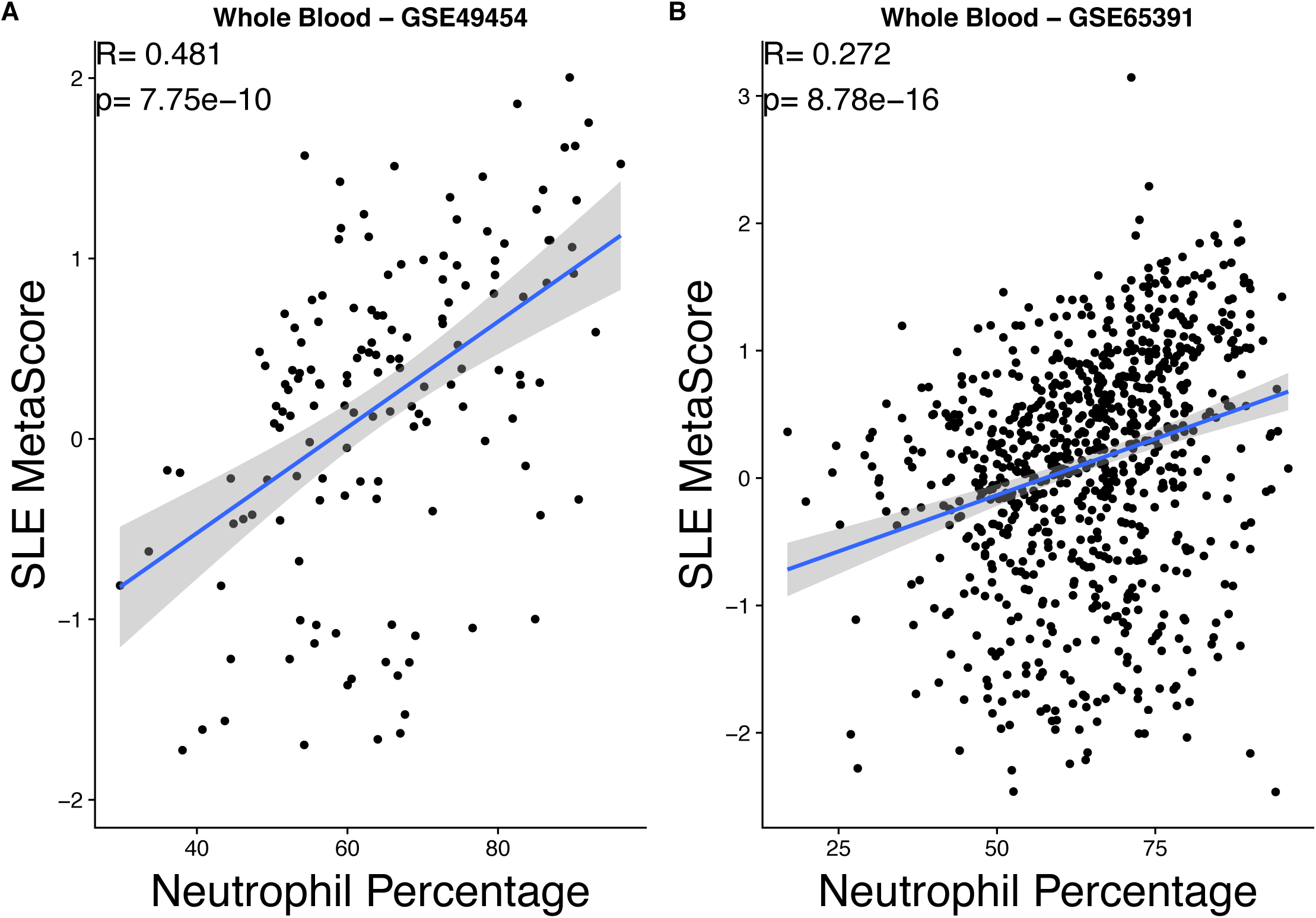
Neutrophil abundance correlation with SLE. In both datasets where neutrophil percentage was provided, we observed a significant correlation between the SLE MetaScore and neutrophil abundance.

**Table S1.**
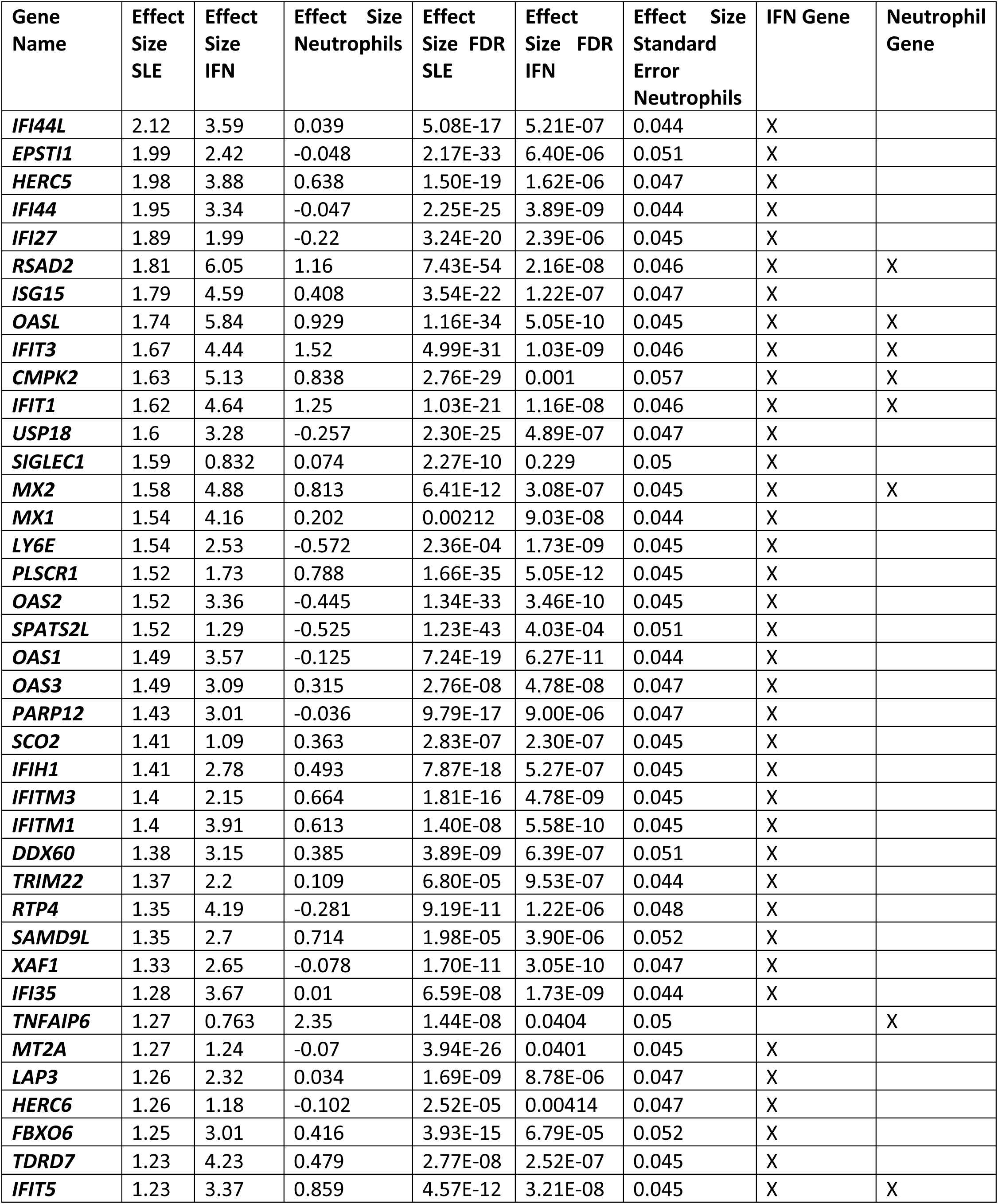
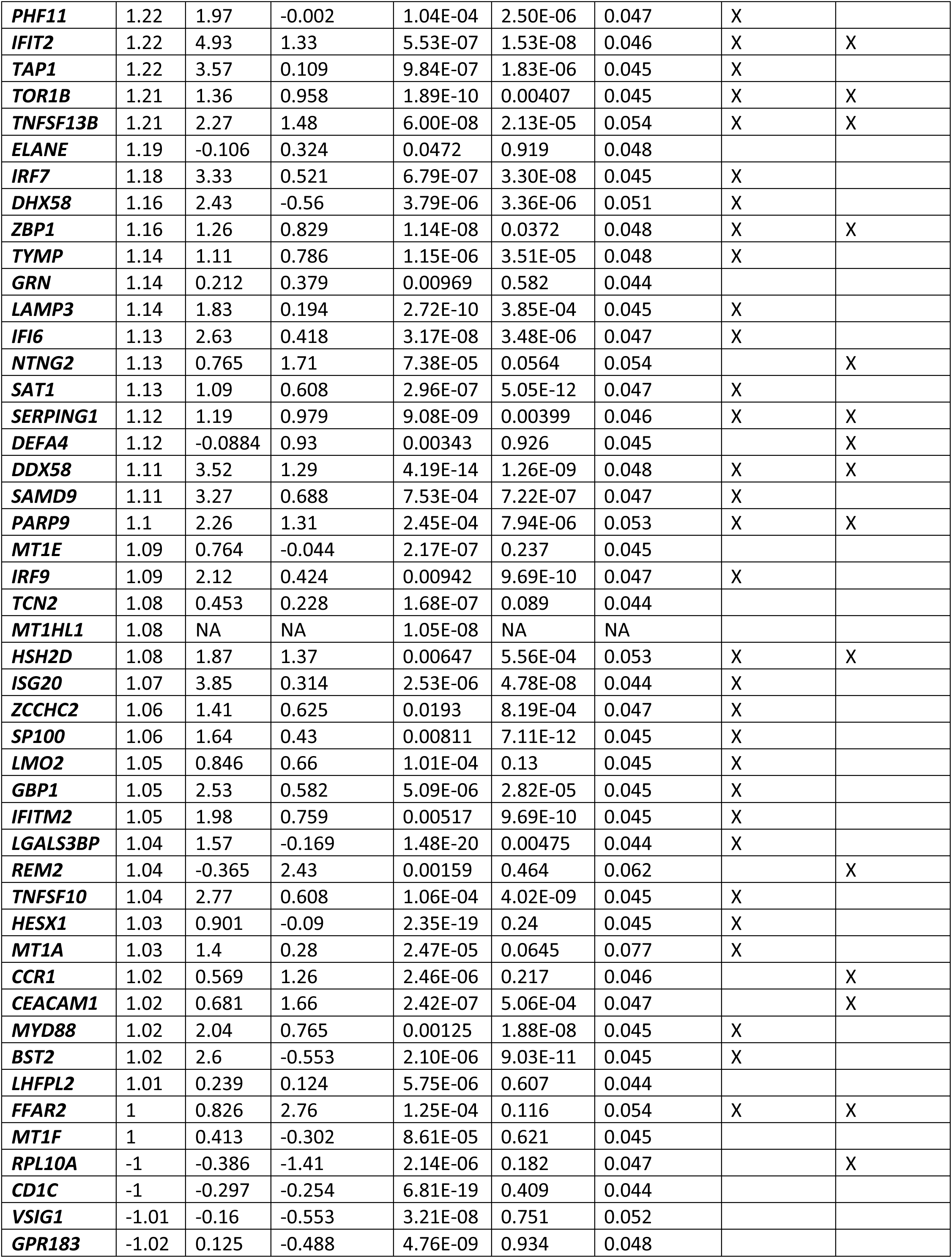
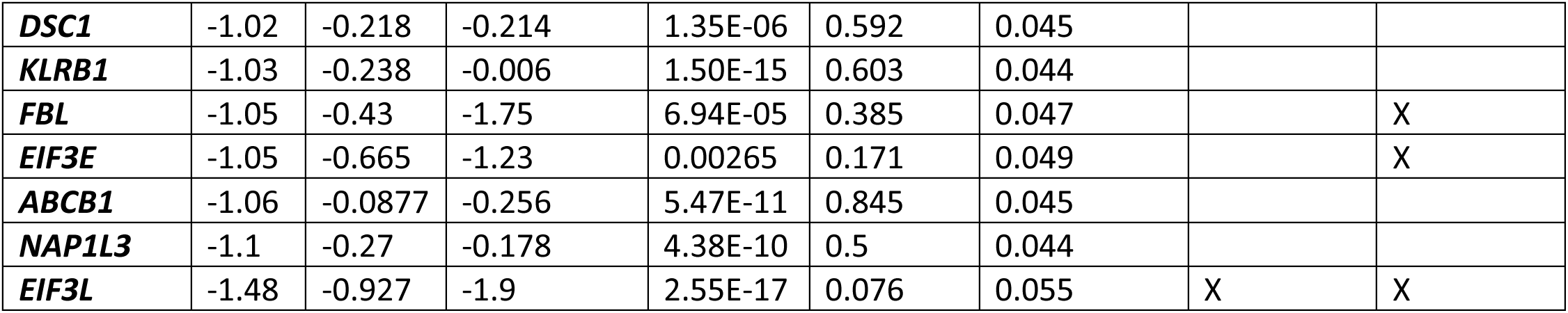
SLE MetaSignature gene names. All genes in the SLE MetaSignature are listed, with their effect sizes and false discovery rates in SLE, IFN, and neutrophils. Additionally, it is indicated whether these genes were considered as IFN or neutrophil genes in our analysis.

| Gene Name | Effect Size SLE | Effect Size IFN | Effect Size Neutrophils | Effect Size FDR SLE | Effect Size FDR IFN | Effect Size Standard Error Neutrophils | IFN Gene | Neutrophil Gene |
| --- | --- | --- | --- | --- | --- | --- | --- | --- |
| *IFI44L* | 2.12 | 3.59 | 0.039 | 5.08E-17 | 5.21E-07 | 0.044 | X |  |
| *EPSTI1* | 1.99 | 2.42 | -0.048 | 2.17E-33 | 6.40E-06 | 0.051 | X |  |
| *HERC5* | 1.98 | 3.88 | 0.638 | 1.50E-19 | 1.62E-06 | 0.047 | X |  |
| *IFI44* | 1.95 | 3.34 | -0.047 | 2.25E-25 | 3.89E-09 | 0.044 | X |  |
| *IFI27* | 1.89 | 1.99 | -0.22 | 3.24E-20 | 2.39E-06 | 0.045 | X |  |
| *RSAD2* | 1.81 | 6.05 | 1.16 | 7.43E-54 | 2.16E-08 | 0.046 | X | X |
| *ISG15* | 1.79 | 4.59 | 0.408 | 3.54E-22 | 1.22E-07 | 0.047 | X |  |
| *OASL* | 1.74 | 5.84 | 0.929 | 1.16E-34 | 5.05E-10 | 0.045 | X | X |
| *IFIT3* | 1.67 | 4.44 | 1.52 | 4.99E-31 | 1.03E-09 | 0.046 | X | X |
| *CMPK2* | 1.63 | 5.13 | 0.838 | 2.76E-29 | 0.001 | 0.057 | X | X |
| *IFIT1* | 1.62 | 4.64 | 1.25 | 1.03E-21 | 1.16E-08 | 0.046 | X | X |
| *USP18* | 1.6 | 3.28 | -0.257 | 2.30E-25 | 4.89E-07 | 0.047 | X |  |
| *SIGLEC1* | 1.59 | 0.832 | 0.074 | 2.27E-10 | 0.229 | 0.05 | X |  |
| *MX2* | 1.58 | 4.88 | 0.813 | 6.41E-12 | 3.08E-07 | 0.045 | X | X |
| *MX1* | 1.54 | 4.16 | 0.202 | 0.00212 | 9.03E-08 | 0.044 | X |  |
| *LY6E* | 1.54 | 2.53 | -0.572 | 2.36E-04 | 1.73E-09 | 0.045 | X |  |
| *PLSCR1* | 1.52 | 1.73 | 0.788 | 1.66E-35 | 5.05E-12 | 0.045 | X |  |
| *OAS2* | 1.52 | 3.36 | -0.445 | 1.34E-33 | 3.46E-10 | 0.045 | X |  |
| *SPATS2L* | 1.52 | 1.29 | -0.525 | 1.23E-43 | 4.03E-04 | 0.051 | X |  |
| *OAS1* | 1.49 | 3.57 | -0.125 | 7.24E-19 | 6.27E-11 | 0.044 | X |  |
| *OAS3* | 1.49 | 3.09 | 0.315 | 2.76E-08 | 4.78E-08 | 0.047 | X |  |
| *PARP12* | 1.43 | 3.01 | -0.036 | 9.79E-17 | 9.00E-06 | 0.047 | X |  |
| *SCO2* | 1.41 | 1.09 | 0.363 | 2.83E-07 | 2.30E-07 | 0.045 | X |  |
| *IFIH1* | 1.41 | 2.78 | 0.493 | 7.87E-18 | 5.27E-07 | 0.045 | X |  |
| *IFITM3* | 1.4 | 2.15 | 0.664 | 1.81E-16 | 4.78E-09 | 0.045 | X |  |
| *IFITM1* | 1.4 | 3.91 | 0.613 | 1.40E-08 | 5.58E-10 | 0.045 | X |  |
| *DDX60* | 1.38 | 3.15 | 0.385 | 3.89E-09 | 6.39E-07 | 0.051 | X |  |
| *TRIM22* | 1.37 | 2.2 | 0.109 | 6.80E-05 | 9.53E-07 | 0.044 | X |  |
| *RTP4* | 1.35 | 4.19 | -0.281 | 9.19E-11 | 1.22E-06 | 0.048 | X |  |
| *SAMD9L* | 1.35 | 2.7 | 0.714 | 1.98E-05 | 3.90E-06 | 0.052 | X |  |
| *XAF1* | 1.33 | 2.65 | -0.078 | 1.70E-11 | 3.05E-10 | 0.047 | X |  |
| *IFI35* | 1.28 | 3.67 | 0.01 | 6.59E-08 | 1.73E-09 | 0.044 | X |  |
| *TNFAIP6* | 1.27 | 0.763 | 2.35 | 1.44E-08 | 0.0404 | 0.05 |  | X |
| *MT2A* | 1.27 | 1.24 | -0.07 | 3.94E-26 | 0.0401 | 0.045 | X |  |
| *LAP3* | 1.26 | 2.32 | 0.034 | 1.69E-09 | 8.78E-06 | 0.047 | X |  |
| *HERC6* | 1.26 | 1.18 | -0.102 | 2.52E-05 | 0.00414 | 0.047 | X |  |
| *FBXO6* | 1.25 | 3.01 | 0.416 | 3.93E-15 | 6.79E-05 | 0.052 | X |  |
| *TDRD7* | 1.23 | 4.23 | 0.479 | 2.77E-08 | 2.52E-07 | 0.045 | X |  |
| *IFIT5* | 1.23 | 3.37 | 0.859 | 4.57E-12 | 3.21E-08 | 0.045 | X | X |
| **PHF11** | 1.22 | 1.97 | -0.002 | 1.04E-04 | 2.50E-06 | 0.047 | X |  |
| **IFIT2** | 1.22 | 4.93 | 1.33 | 5.53E-07 | 1.53E-08 | 0.046 | X | X |
| **TAP1** | 1.22 | 3.57 | 0.109 | 9.84E-07 | 1.83E-06 | 0.045 | X |  |
| **TOR1B** | 1.21 | 1.36 | 0.958 | 1.89E-10 | 0.00407 | 0.045 | X | X |
| **TNFSF13B** | 1.21 | 2.27 | 1.48 | 6.00E-08 | 2.13E-05 | 0.054 | X | X |
| **ELANE** | 1.19 | -0.106 | 0.324 | 0.0472 | 0.919 | 0.048 |  |  |
| **IRF7** | 1.18 | 3.33 | 0.521 | 6.79E-07 | 3.30E-08 | 0.045 | X |  |
| **DHX58** | 1.16 | 2.43 | -0.56 | 3.79E-06 | 3.36E-06 | 0.051 | X |  |
| **ZBP1** | 1.16 | 1.26 | 0.829 | 1.14E-08 | 0.0372 | 0.048 | X | X |
| **TYMP** | 1.14 | 1.11 | 0.786 | 1.15E-06 | 3.51E-05 | 0.048 | X |  |
| **GRN** | 1.14 | 0.212 | 0.379 | 0.00969 | 0.582 | 0.044 |  |  |
| **LAMP3** | 1.14 | 1.83 | 0.194 | 2.72E-10 | 3.85E-04 | 0.045 | X |  |
| **IFI6** | 1.13 | 2.63 | 0.418 | 3.17E-08 | 3.48E-06 | 0.047 | X |  |
| **NTNG2** | 1.13 | 0.765 | 1.71 | 7.38E-05 | 0.0564 | 0.054 |  | X |
| **SAT1** | 1.13 | 1.09 | 0.608 | 2.96E-07 | 5.05E-12 | 0.047 | X |  |
| **SERPING1** | 1.12 | 1.19 | 0.979 | 9.08E-09 | 0.00399 | 0.046 | X | X |
| **DEFA4** | 1.12 | -0.0884 | 0.93 | 0.00343 | 0.926 | 0.045 |  | X |
| **DDX58** | 1.11 | 3.52 | 1.29 | 4.19E-14 | 1.26E-09 | 0.048 | X | X |
| **SAMD9** | 1.11 | 3.27 | 0.688 | 7.53E-04 | 7.22E-07 | 0.047 | X |  |
| **PARP9** | 1.1 | 2.26 | 1.31 | 2.45E-04 | 7.94E-06 | 0.053 | X | X |
| **MT1E** | 1.09 | 0.764 | -0.044 | 2.17E-07 | 0.237 | 0.045 |  |  |
| **IRF9** | 1.09 | 2.12 | 0.424 | 0.00942 | 9.69E-10 | 0.047 | X |  |
| **TCN2** | 1.08 | 0.453 | 0.228 | 1.68E-07 | 0.089 | 0.044 |  |  |
| **MT1HL1** | 1.08 | NA | NA | 1.05E-08 | NA | NA |  |  |
| **HSH2D** | 1.08 | 1.87 | 1.37 | 0.00647 | 5.56E-04 | 0.053 | X | X |
| **ISG20** | 1.07 | 3.85 | 0.314 | 2.53E-06 | 4.78E-08 | 0.044 | X |  |
| **ZCCHC2** | 1.06 | 1.41 | 0.625 | 0.0193 | 8.19E-04 | 0.047 | X |  |
| **SP100** | 1.06 | 1.64 | 0.43 | 0.00811 | 7.11E-12 | 0.045 | X |  |
| **LMO2** | 1.05 | 0.846 | 0.66 | 1.01E-04 | 0.13 | 0.045 | X |  |
| **GBP1** | 1.05 | 2.53 | 0.582 | 5.09E-06 | 2.82E-05 | 0.045 | X |  |
| **IFITM2** | 1.05 | 1.98 | 0.759 | 0.00517 | 9.69E-10 | 0.045 | X |  |
| **LGALS3BP** | 1.04 | 1.57 | -0.169 | 1.48E-20 | 0.00475 | 0.044 | X |  |
| **REM2** | 1.04 | -0.365 | 2.43 | 0.00159 | 0.464 | 0.062 |  | X |
| **TNFSF10** | 1.04 | 2.77 | 0.608 | 1.06E-04 | 4.02E-09 | 0.045 | X |  |
| **HESX1** | 1.03 | 0.901 | -0.09 | 2.35E-19 | 0.24 | 0.045 | X |  |
| **MT1A** | 1.03 | 1.4 | 0.28 | 2.47E-05 | 0.0645 | 0.077 | X |  |
| **CCR1** | 1.02 | 0.569 | 1.26 | 2.46E-06 | 0.217 | 0.046 |  | X |
| **CEACAM1** | 1.02 | 0.681 | 1.66 | 2.42E-07 | 5.06E-04 | 0.047 |  | X |
| **MYD88** | 1.02 | 2.04 | 0.765 | 0.00125 | 1.88E-08 | 0.045 | X |  |
| **BST2** | 1.02 | 2.6 | -0.553 | 2.10E-06 | 9.03E-11 | 0.045 | X |  |
| **LHFPL2** | 1.01 | 0.239 | 0.124 | 5.75E-06 | 0.607 | 0.044 |  |  |
| **FFAR2** | 1 | 0.826 | 2.76 | 1.25E-04 | 0.116 | 0.054 | X | X |
| **MT1F** | 1 | 0.413 | -0.302 | 8.61E-05 | 0.621 | 0.045 |  |  |
| **RPL10A** | -1 | -0.386 | -1.41 | 2.14E-06 | 0.182 | 0.047 |  | X |
| **CD1C** | -1 | -0.297 | -0.254 | 6.81E-19 | 0.409 | 0.044 |  |  |
| **VSIG1** | -1.01 | -0.16 | -0.553 | 3.21E-08 | 0.751 | 0.052 |  |  |
| **GPR183** | -1.02 | 0.125 | -0.488 | 4.76E-09 | 0.934 | 0.048 |  |  |
| ***DSC1*** | -1.02 | -0.218 | -0.214 | 1.35E-06 | 0.592 | 0.045 |  |  |
| ***KLRB1*** | -1.03 | -0.238 | -0.006 | 1.50E-15 | 0.603 | 0.044 |  |  |
| ***FBL*** | -1.05 | -0.43 | -1.75 | 6.94E-05 | 0.385 | 0.047 |  | X |
| ***EIF3E*** | -1.05 | -0.665 | -1.23 | 0.00265 | 0.171 | 0.049 |  | X |
| ***ABCB1*** | -1.06 | -0.0877 | -0.256 | 5.47E-11 | 0.845 | 0.045 |  |  |
| ***NAP1L3*** | -1.1 | -0.27 | -0.178 | 4.38E-10 | 0.5 | 0.044 |  |  |
| ***EIF3L*** | -1.48 | -0.927 | -1.9 | 2.55E-17 | 0.076 | 0.055 | X | X |

**Table S2.** Interferon datasets. Data downloaded from GEO.

| Dataset | Center | Cell Type | IFN Type | Stim Length | Sample Size | Reference |
| --- | --- | --- | --- | --- | --- | --- |
| GSE14386 | University of North Carolina, Chapel Hill, NC | PBMCs | IFNβ -1a | 24h | 30 | (168) |
| GSE16755 | NIH, Bethesda, ND | Monocyte derived macrophages | IFNα | 1-4h | 6 | (169) |
| GSE17301 | Center for Applied Medical Research, Pamplona, Spain | CD8+CD45R0- T cells | IFNα-2b or IFNα-5 | 48h | 15 | (170) |
| GSE1740 | The Hospital for Special Surgery, New York, NY | Macrophages | IFNα | 3h | 12 | (171) |
| GSE23307 | Cleveland Clinic, Cleveland, OH | Whole blood | IFNβ | 3h | 6 | (172) |
| GSE27337 | University of Texas Southwestern, Dallas, TX | CD8+CD45RA+ T cells | IFNα | 84h | 48 | (173) |
| GSE30536 | Universite Laval, Quebec, Canada | Macrophages | IFNα-2 | 18h | 17 | (174) |
| GSE34627 | Technion Institute, Haifa, Israel | Monocytes and T cells | IFNβ | 16h | 12 | (175) |
| GSE36287 | Harvard Universty, Boston, MA | Keratinocytes | IFNα | | 24 | (176) |
| GSE37715 | Hiroshima University, Hiroshima, Japan | Hepatocytes | IFNα | 6h | 15 | |
| GSE38147 | University of Basel, Basel, Switzerland | Hepatocytes | IFNα | 6-24h | 10 | (177) |
| GSE38351 | Deutsches Rheuma-Forschungszentrum, Berlin, Germany | Monocytes | IFNα-2a | 1.5h | 74 | (152) |
| GSE3920 | Medical School of Turin, Turin, Italy | Endothelial cells and fibroblasts | IFNα, IFNβ | 5h | 23 | (178) |
| GSE47616 | Northwestern University, Chicago, IL | Fibroblasts | IFNα, IFNβ | 24h | 22 | (179) |
| GSE7509 | Yale University, New Haven, CT | Monocyte derived dendritic cells | IFNα | 24h | 26 | (180) |

**Table S3.** Neutrophil, NK cell, and heavy metal datasets. Data downloaded from GEO.

| **Data Type** | **Dataset** | **Center** | **Sample Size** | **Reference** |
| --- | --- | --- | --- | --- |
| NETosis Stimulations | GSE80489 | All India Institute of Medical Science, New Delhi, India | 18 | (181) |
| Neutropoeisis | GSE42519 | Copenhagen University, Copenhagen, Denmark | 34 | (182) |
| Neutropoeisis | GSE19556 | Princeton University, Princeton, NJ | 24 | (183) |
| Neutropoeisis | GSE49883 | Karolinska Institute, Stockholm, Sweden | 20 | (184) |
| Cadmium Exposure of MCF7 Cell Line | GSE52404 | University of Skovde, Skovde, Sweden | 7 |  |
| Zinc Exposure of A549 Lung Cells | GSE6972 | University of Southern California, Los Angeles, CA | 24 | (42) |

**Table S4.**
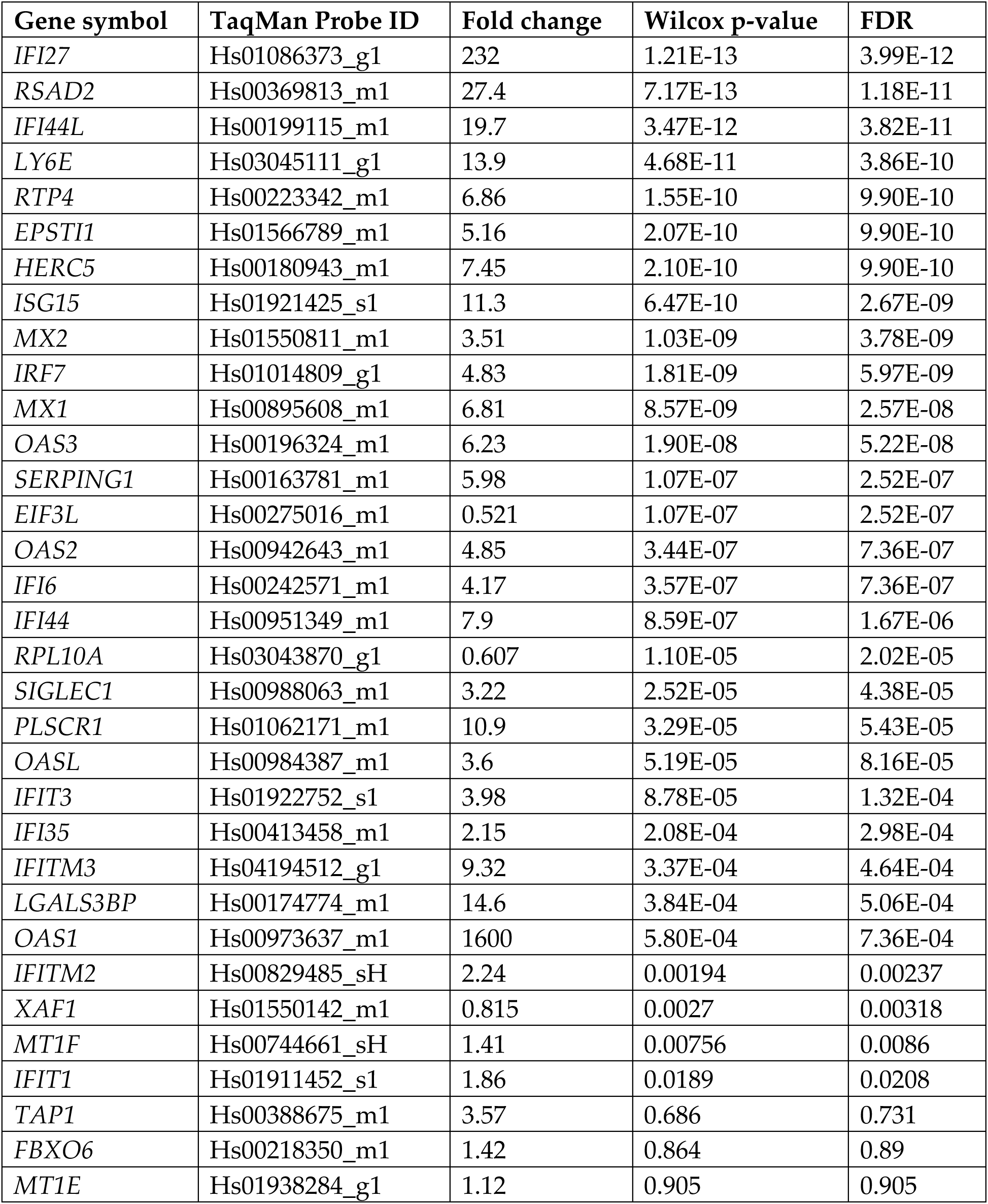
Gene-level prospective validation results. Gene name, probe ID, fold change, p-value, and FDR for all genes from the SLE MetaSignature which were run on the prospective Fluidigm RT-qPCR assay.

| Gene symbol | TaqMan Probe ID | Fold change | Wilcox p-value | FDR |
| --- | --- | --- | --- | --- |
| *IFI27* | Hs01086373_g1 | 232 | 1.21E-13 | 3.99E-12 |
| *RSAD2* | Hs00369813_m1 | 27.4 | 7.17E-13 | 1.18E-11 |
| *IFI44L* | Hs00199115_m1 | 19.7 | 3.47E-12 | 3.82E-11 |
| *LY6E* | Hs03045111_g1 | 13.9 | 4.68E-11 | 3.86E-10 |
| *RTP4* | Hs00223342_m1 | 6.86 | 1.55E-10 | 9.90E-10 |
| *EPSTI1* | Hs01566789_m1 | 5.16 | 2.07E-10 | 9.90E-10 |
| *HERC5* | Hs00180943_m1 | 7.45 | 2.10E-10 | 9.90E-10 |
| *ISG15* | Hs01921425_s1 | 11.3 | 6.47E-10 | 2.67E-09 |
| *MX2* | Hs01550811_m1 | 3.51 | 1.03E-09 | 3.78E-09 |
| *IRF7* | Hs01014809_g1 | 4.83 | 1.81E-09 | 5.97E-09 |
| *MX1* | Hs00895608_m1 | 6.81 | 8.57E-09 | 2.57E-08 |
| *OAS3* | Hs00196324_m1 | 6.23 | 1.90E-08 | 5.22E-08 |
| *SERPING1* | Hs00163781_m1 | 5.98 | 1.07E-07 | 2.52E-07 |
| *EIF3L* | Hs00275016_m1 | 0.521 | 1.07E-07 | 2.52E-07 |
| *OAS2* | Hs00942643_m1 | 4.85 | 3.44E-07 | 7.36E-07 |
| *IFI6* | Hs00242571_m1 | 4.17 | 3.57E-07 | 7.36E-07 |
| *IFI44* | Hs00951349_m1 | 7.9 | 8.59E-07 | 1.67E-06 |
| *RPL10A* | Hs03043870_g1 | 0.607 | 1.10E-05 | 2.02E-05 |
| *SIGLEC1* | Hs00988063_m1 | 3.22 | 2.52E-05 | 4.38E-05 |
| *PLSCR1* | Hs01062171_m1 | 10.9 | 3.29E-05 | 5.43E-05 |
| *OASL* | Hs00984387_m1 | 3.6 | 5.19E-05 | 8.16E-05 |
| *IFIT3* | Hs01922752_s1 | 3.98 | 8.78E-05 | 1.32E-04 |
| *IFI35* | Hs00413458_m1 | 2.15 | 2.08E-04 | 2.98E-04 |
| *IFITM3* | Hs04194512_g1 | 9.32 | 3.37E-04 | 4.64E-04 |
| *LGALS3BP* | Hs00174774_m1 | 14.6 | 3.84E-04 | 5.06E-04 |
| *OAS1* | Hs00973637_m1 | 1600 | 5.80E-04 | 7.36E-04 |
| *IFITM2* | Hs00829485_sH | 2.24 | 0.00194 | 0.00237 |
| *XAF1* | Hs01550142_m1 | 0.815 | 0.0027 | 0.00318 |
| *MT1F* | Hs00744661_sH | 1.41 | 0.00756 | 0.0086 |
| *IFIT1* | Hs01911452_s1 | 1.86 | 0.0189 | 0.0208 |
| *TAP1* | Hs00388675_m1 | 3.57 | 0.686 | 0.731 |
| *FBXO6* | Hs00218350_m1 | 1.42 | 0.864 | 0.89 |
| *MT1E* | Hs01938284_g1 | 1.12 | 0.905 | 0.905 |

**Table S5.** Disease activity correlations of all SLE MetaScore and subsets. Correlation of SLE MetaScore or subset with the patient SLEDAI values. Partial data is presented in Figure 6C.

| **Dataset** | **SLE**<br>**MetaSignature** | **IFN SLE**<br>**MetaSignature** | **Non-IFN SLE**<br>**MetaSignature** | **Neutrophil SLE**<br>**MetaSignature** | **Under-**<br>**appreciated**<br>**SLE**<br>**MetaSignature** |
| --- | --- | --- | --- | --- | --- |
| Neutrophils-GSE27427 | 0.284 | 0.254 | 0.354 | 0.387 | 0.197 |
| PBMCs- GSE8650 | 0.67 | 0.545 | 0.782 | 0.77 | 0.712 |
| Whole Blood-GSE49454 | 0.157 | 0.113 | 0.247 | 0.197 | 0.283 |
| Whole Blood-GSE65391 | 0.281 | 0.252 | 0.342 | 0.295 | 0.339 |
| PBMCs- GSE88884 | 0.25 | 0.236 | 0.233 | 0.193 | 0.243 |

## Supplement S1. Dataset descriptions

### Discovery datasets

**GSE17755** (22 SLE, 22 healthy; 44 total)

Lee et al. profiled peripheral blood samples from 21 patients (all women, median age: 35 years, range: 26-72 years) with SLE according to the diagnostic criteria of the American College of Rheumatology (ACR) and 45 healthy individuals (23 males, 22 females). Because all patients with SLE were women, we only used healthy females as controls in this cohort, and discarded data from healthy males.

**GSE8650:** (38 SLE, 21 healthy; 59 total)

Allantaz et al. profiled PBMC samples from 19 pediatric patients with system juvenile idiopathic arthritis (sJIA) during the systemic phase of the disease (fever and/or arthritis), 25 sJIA patients with no systemic symptoms (arthritis only or no symptoms), 39 healthy controls, 94 pediatric patients with acute viral and bacterial infections (available under GSE6269), 38 pediatric patients with Systemic Lupus Erythematosus (SLE), and 6 patients with a second IL-1 mediated disease known as PAPA syndrome. We used transcriptome data only from healthy controls and pediatric patients with SLE in the discovery analysis. Rest of the samples were used in downstream analysis after the signature for SLE was derived.

**GSE50635** (33 SLE, 16 healthy; 49 total)

Ko *et al.* profiled whole blood samples from 33 female patients with SLE and 16 matched controls from European-American (EA) and African-American (AA) ancestral backgrounds. This dataset is not associated with any publication in the NCBI GEO database.

**GSE39088** (26 SLE, 34 healthy; 60 total)

Lauwerys *et al.* profiled 28 female patients with SLE (aged 18–50 years; recruited in a multicenter, randomized, double-blind placebo-controlled, phase I/II staggered dose-escalation trial of IFN-K (ClinicalTrials.gov registry number <u>NCT01058343</u>). Patients were randomized to receive three or four injections of placebo (n = 7) or 30 µg (n = 3), 60 µg (n = 6), 120 µg (n = 6) or 240 µg (n = 6) IFN-K. We only used samples prior to treatment initiation in the analysis.

**GSE22098 (**28 adult and 12 pediatric SLE, 42 healthy; 83 total)

Berry et al. collected whole blood sample from 12 pediatric streptococcus, 40 pediatric staphylococcus, 31 still’s disease, 82 pediatric systemic lupus erythematosus (SLE) and 28 adult SLE patients. RNA was extracted and globin reduced. Labeled cRNA was hybridized to Illumina Human HT-12 Beadchips. Healthy controls were included to match patients’ demographic data. Genespring software was used to analyze active TB transcript signatures, comparing with healthy controls and other inflammatory and infectious diseases.

**GSE11909:** (63 SLE, 12 healthy; 75 total) First visit patient data

Chaussabel et al. profiled 239 PBMC samples from individuals with one of the following conditions: systemic juvenile idiopathic arthritis (n=47), systemic lupus erythematosus (n=63), type I diabetes (n=20), metastatic melanoma (n=39), acute infections (*Escherichia coli* (n=22), *Staphylococcus aureus* (n=18), Influenza A (n=16)), or liver transplant recipients undergoing immunosuppressive therapy (n=37). We only used data only from the first visit.

### Validation datasets

**GSE12374 ()**

: Lee et al. obtained peripheral blood from female SLE patients (n = 11 median age 35 years, range 27 to 72 years) and healthy women (n=6). Gene expression profile was analyzed using DNA microarray covering 30,000 human genes. Differentially expressed immune response-related genes were selected and analyzed by using Expression Analysis Systemic Explorer (EASE) based on Gene Ontology (GO) followed by network pathway analysis with Ingenuity Pathways Analysis (IPA).

**GSE24706:** Li et al. obtained 10 healthy control (HC) samples with high ANA, 7 first degree relative of SLE patient (FDR) with high ANA, 10 HCs with low ANA, 6 FDR with low ANA, 15 SLE patients (SLE).

The overall study group included 1,159 individuals from DRADR: 401 healthy controls (HC) who were negative for current or past autoimmune disease, 116 first-degree relatives (FDR), 294 patients with SLE, 151 patients with less than 4 SLE criteria and considered as having incomplete lupus (ILE), 154 with rheumatoid arthritis (RA) and 43 with other miscellaneous conditions including scleroderma, Sjogren’s syndrome, ankylosing spondylitis and vasculitis. More detailed analyses were carried out on a subset of HC individuals with high ANA values (*n* = 18) and these were compared to gender- and age-matched HC with negative ANA values (*n* = 16) and to SLE patients with high ANA levels of >100 E.U. (*n* = 14).

Measurements carried out on serum samples included ANA, extractable nuclear antibodies (ENA) and autoantibody profiling using an array with more than 100 specificities. Whole blood RNA samples from a subset of individuals were used to analyze gene expression on the Illumina platform. Data were analyzed for associations of high ANA levels with demographic features, the presence of other autoantibodies and with gene expression profiles.

**GSE49454:** Chiche el al. enrolled sixty-two consecutive patients with SLE fulfilling the 1997 ACR criteria were enrolled between 2009 and 2011 in the Departments of Internal Medicine and Nephrology at a French reference center for autoimmune diseases (Hôpital de la Conception, Marseille, France) and followed-up them prospectively.

SLE patients were split into three groups (Table S1). The “at inclusion” group included all SLE patients at their first visit, irrespective of SLE disease activity at that time. The “quiescent” group included SLE patients at their first available visit with low disease activity, defined by no flare or treatment modifications for at least 60 days prior to the visit, and a SLEDAI of ≤4. The “longitudinal” group included SLE patients who had at least three consecutive visits during the study.

**GSE61635:** The goal of this study was to characterize gene expression profiles in RNP autoantibody+ SLE versus healthy blood donors with a focus on select cytokines that may be important in B cell activation and differentiation, including BAFF, IL-21, and IL-33. Affymetrix microarrays were used to characterize the global program of gene expression in the SLE patients, and to identify differentially expressed genes in patients compared to healthy controls. mRNA from the blood of a SLE cohort (79 patients with some repeat visits for a total of 99 arrays) and 30 healthy volunteers (one array per volunteer) were analyzed.

There were 73 female and 6 male subjects. Disease duration ranged from 0 to 453 months with a median of 37.5 months. SLE Disease Activity Index (SLEDAI) ranged from 0 to 31 with a median of 6.

**GSE65391:** Banchereau R et al. longitudinally profiled the whole blood transcriptomes of 158 SLE patients by microarray for up to 4 years, yielding 924 SLE samples and 48 matched pediatric healthy samples. The transcriptional data are complemented by demographic, laboratory and clinical data. They confirmed a prevalent IFN signature and identified a plasmablast signature as the most robust biomarker of DA.They also detected gradual enrichment of neutrophil transcripts during progression to active nephritis, and distinct signatures in response to treatment in different nephritis subclasses.

**GSE72798:** This cohort was generated to validate if IFNalpha kinoid induces neutralizing anti-IFNalpha antibodies that decrease the expression of IFN-induced and B cell activation associated transcripts: analysis of extended follow-up data from the IFN-K phase I/II study

**Cohort has 82 total samples with 10 healthy and rest SLE, we have only included patient data for first visit for SLE patients.**

**GSE81622:** This cohort was generated to perform whole genome transcription and DNA methylation analysis in PBMC of 30 SLE patients, including 15 with LN (SLE LN+) and 15 without LN (SLE LN-), and 25 normal controls (NC) using HumanHT-12 Beadchips and Illumina Human Methy450 chips. The serum pro-inflammatory cytokines were quantified using Bio-plex human cytokine 27-plex assay.

### Extended validation

**GSE10325 :** Becker et al. enrolled SLE patients fulfilled at least 4 of 11 American College of Rheumatology classification criteria for SLE. Disease activity assessed at the time of blood acquisition was calculated using the systemic lupus erythematosus disease activity index (SLEDAI). They compared PBMC subsets from a total of fifteen female SLE patients (mean age 39±12 years) and eleven female HC (mean age 37±10 years). Although patients were on a variety of disease modifying agents, patients on high dose immunocytotoxic therapies or steroids were excluded from the study. However, patients on lower doses of prednisone (10–20 mg/day; and 1 patient on 40 mg/day) were included.

**GSE13887:** Fernandez et al. investigated a total of 44 Caucasian female patients with systemic lupus erythematosus (SLE) in their cohort. Disease activity was assessed by the SLEDAI score^72^. Six patients were treated with rapamycin 2 mg/day (age: 40 ± 8.3 years; SLEDAI: 0.8). Among the 38 remaining SLE patients treated without rapamycin, 28 were receiving prednisone (5–50 mg/day) and immunosupppresive drugs including hydroxychloroquine (400 mg/day), mycophenolate mofetil (3 g/day), cyclosporin A (50–100 mg/day). Their mean age was 36.3 ± 4.3 years, ranging between 18–60; SLEDAI: 1.3 ± 0.9. Furthermore, ten patients (age: 38.5 ± 6.4) SLEDAI: 4.8 ± 3.8) were freshly diagnosed and had not been treated with prednisone or cytotoxic drugs.

These patients and five additional patients that have received prednisone or cytotoxic drugs provided cells for microarray analysis. As controls, 23 age-matched healthy female subjects and 8 female patients with rheumatoid arthritis (RA; age: 51.3 ± 6.7 years) ^73^ were studied. RA patients were treated with methotrexate, cyclosporin A, leflunomide, etanercept, or adalimumab. The study has been approved by the Institutional Review Board for the Protection of Human Subjects.

**GSE24060:** O’Hanlon et al. enrolled five adult (at least 18 years of age) and 15 juvenile MZ twin pairs discordant for SAID and 40 unrelated control subjects (two controls per twin pair) matched on age within 6 years, gender, and ethnicity in this study. These subjects were enrolled between 2001 and 2006 in the National Institutes of Health (NIH) investigational review board-approved Twins-Sib study assessing the pathogenesis of SAID. Twin pairs enrolled within 4 years of probands’ diagnoses included 19 non-Hispanic Caucasian twin pairs and a single Hispanic twin pair (with SLE). Probands fulfilled American College of Rheumatology criteria for adult or juvenile SLE (*n* = 4 and 2, respectively), RA or JRA (*n* = 1 and 5, respectively), juvenile dermatomyositis (JDM) (*n* = 7), or juvenile polymyositis (JPM) (*n* = 1); they excluded patients with inherited, metabolic, infectious, or other causes of disease. The juvenile probands ranged in age from 3 to 18 years (mean of 11.2 years), whereas adults ranged from 19 to 43 years (mean of 29.2 years). Twins included 14 female and 6 male pairs. Monozygosity was confirmed by short tandem repeat analysis of genomic DNAs (Proactive Genetics, Inc., Augusta, GA, USA). Unrelated, matched controls were free of infections, trauma, vaccines, and surgeries for 8 weeks and had no first-degree family members with SAID.

**GSE26949:** Thacker et al. enrolled the patients and obtained peripheral blood, from the university of Michigan outpatient Rheumatology clinic who fulfilled the revised American College of Rheumatology criteria for SLE. Age- and gender-matched healthy controls were recruited by advertisement. Lupus disease activity was assessed by the SLE Disease Activity Index (SLEDAI). In overall experiment, Human healthy EPCs and CACs from PBMCs were isolated and cultured under proangiogenic stimulation; after IFNa incubation or not, RNA was extracted and processed for hybridization on Affymetrix microarrays.

**GSE26950:** Thacker et al. enrolled the patients and obtained peripheral blood, from the university of Michigan outpatient Rheumatology clinic who fulfilled the revised American College of Rheumatology criteria for SLE. Age- and gender-matched healthy controls were recruited by advertisement. Lupus disease activity was assessed by the SLE Disease Activity Index (SLEDAI). In overall experiment, Human lupus EPCs and CACs from PBMCs were isolated and cultured under proangiogenic stimulation; after IFNa incubation or not, RNA was extracted and processed for hybridization on Affymetrix microarrays.

**GSE26975:** Villanueva et al. enrolled Lupus patients from the University of Michigan outpatient rheumatology clinic who fulfilled the revised American College of Rheumatology criteria for SLE. Disease activity was assessed by the SLE disease activity index. Gender-matched healthy controls were recruited by advertisement. Demographic and clinical information on the lupus patients. In overall experiment they isolated neutrophils and LDGs from PBMCs. RNA from 9 healthy neutrophils, 10 lupus neutrophils and 10 lupus LDGs was extracted and processed for hybridization on Affymetrix microarrays. Lupus patients SLEDAI score was between 0 and 20. Most of the patients were on antimalarials, PDN or on MMF drugs.

**GSE27427:** Garcia-Romo et al. obatained blood samples from patients fulfilling the diagnosis of SLE according to the criteria established by the American College of Rheumatology. Disease activity was assessed by the SLEDAI as measured on the day of blood collection. Healthy pediatric controls were children visiting the clinic either for reasons not related to autoimmunity or infectious diseases or for surgery not associated with any inflammatory diseases. They ran to experiments, (Expt 1) Neutrophils from 21 SLE samples (19 patients) and 12 healthy donors were isolated, and extracted RNAs were used generate microarray data. (Expt 2) Neutrophils isolated from 2 healthy children (not used in the first experiment) were cultured with autologous sera (control), Interferon alpha (100U and 1000U), and 4 SLE sera and 6 SLE sera for 6 hours and RNAs were extract for microarray experiment.

**GSE29536:** This dataset was used to establish whole blood transcriptional modules (n=260) that represent groups of coordinately expressed transcripts that exhibit altered abundance within individual datasets or across multiple datasets. This modular framework was generated to reduce the dimensionality of whole blood microarray data processed on the Illumina Beadchip platform yielding data-driven transcriptional modules with biologic meaning.

This series combines nine independent datasets representing a spectrum of human pathologies expected to result in changes in gene abundance related to changes in expression or cellular composition of whole blood. These nine datasets are composed of 410 individual whole blood profiles generated from patients with HIV, tuberculosis, sepsis, systemic lupus erythematosus, systemic arthritis, B-cell deficiency and liver transplant. For each dataset healthy controls are also included. Each dataset’s expression data was preprocessed independently.

**GSE30153:** Garaud et al. selected 17 patients (15 females and 2 males) ageing from 23 to 59 with the diagnosis of SLE for the study. The SLE diagnosis was based on the presence of at least 4 criterias among those defined by the American College of Rheumatology. The lupus was inactive in these patients for more than 6 months, with a Systemic Lupus Erythematosous Disease Activity Index (SLEDAI) score less than 4, and they did not receive any immunosuppressive drug. If they needed steroids, the patients were not treated with more than 10 mg of prednisone per day (4 patients). 10 patients were treated with hydroxychoroquine. The 10 control subjects were healthy individuals, (8 females and 2 males) ageing from 23 to 53 years, with no personal nor familial history of autoimmune disease.

They compared the peripheral B cell transcriptomes of quiescent lupus patients to normal B cell transcriptomes in this cohort.

**GSE32591:** Berthier et al. collected renal a total of 47 samples from the European Renal cDNA Bank (ERCB), they were processed and used for microarray analysis: 15 pre-transplant healthy living donors (LD) and 32(25 female/7 male) lupus nephritis (LN) patients median age was. For real-time PCR, 11 LD and 9 LN samples were used from an independent cohort (of the ERCB).

RNA from glomeruli and tubulointerstitial compartments was extracted and processed for hybridization on Affymetrix microarrays.

**GSE36700:** Nzeusseu et al. obtained synovial biopsy tissue (15–20 synovial samples per patient) by needle arthroscopy of the affected knee of patients with SLE (n = 6), patients with RA (n = 7), and patients with OA (n = 6).

All patients with SLE met the American College of Rheumatology (ACR; formerly, the American Rheumatism Association) revised classification criteria for SLE, all were female, and the mean age was 32 years (range 19–40 years). All SLE patients had active articular disease at the time of synovial tissue sampling, and none had received immunosuppressive therapy; some of the SLE patients were receiving nonsteroidal antiinflammatory drugs. All patients with RA met the ACR classification criteria for RA and all had early (<1 year’s duration) active disease at the time of tissue sampling. Among the patients with RA, 2 were female and 5 were male, and the mean age was 51 years (range 37–69 years). In these patients, the mean C-reactive protein (CRP) level was 25 mg/liter (range 9–96 mg/liter), and the mean Disease Activity Score in 28 joints (including the CRP) was 5.08 (range 3.76–5.82). None of the RA patients had received any treatment, except with nonsteroidal antiinflammatory drugs. Among the patients with OA, 5 were female and 1 was male, and the mean age was 63.2 years (range 51–73 years).

All patients had a swollen knee at the time of the needle arthroscopy procedure. The biopsy samples were harvested before initiation of disease-modifying antirheumatic drugs or any other immunosuppressive therapy. All of the RA patients were subsequently treated with methotrexate. Tumor necrosis factor (TNF) blocking agents were added to the regimen of 5 of these patients at a later stage. All of the SLE patients subsequently received antimalarial drugs. Combination therapy with methotrexate was later started because of persistent joint involvement in 3 of these patients. Azathioprine was started for severe hematologic manifestations in 2 other patients.

**GSE36941:Terrier et al.** evaluated the safety and the immunological effects of vitamin D supplementation in SLE patients with hypovitaminosis D using transcriptomic study at M0 and M2.

They assessed 24 SLE patients for eligibility (twenty-two women and two men, mean age ± SD, 31 ± 8 years). Their serum 25(OH)D level was measured. Hypovitaminosis D was defined as serum 25(OH)D < 30 ng/mL, while vitamin D sufficiency was defined as serum levels between 30 and 100 ng/mL [17]. Those with hypovitaminosis D (< 30 ng/mL) were placed on the following schedule of oral vitamin D supplementation: 100,000 IU of cholecalciferol per week for 4 weeks, followed by 100,000 IU of cholecalciferol per month for 6 months. All supplemented patients were screened before vitamin D supplementation (Day 0, or D0), and 2 and 6 months (M2 and M6) after the beginning of vitamin D supplementation. All but four patients received hydroxychloroquine (200 or 400 mg daily) and/or oral prednisone (≤ 15 mg/day, median dosage 5 mg/day). Three patients received a stable dosage of immunosuppressive agents. The study was approved by the institutional ethics committee, the Comité de protection des personnes Ile-de-France VI, in the Pitié-Salpêtrière Hospital (Paris, France) and informed consent was obtained from all patients.

**GSE37356:** In this cohort, monocytes were obtained from 20 patients with SLE and 16 healthy controls and were in vitro differentiated into macrophages. Subjects also underwent laboratory and imaging studies of the coronary arteries, carotid arteries, and aorta to evaluate for subclinical atherosclerosis.

**GSE37573:** In this cohort, Epstein-Barr virus (EBV) transformed B cells derived from two patients with systemic lupus erythematosus (SLE) and two normal unrelated controls were stimulated with a biologically relevant signal, co-crosslinking of the B cell antigen receptor (BCR) and FcγR2b. Total RNA was isolated at various timepoints post-stimulation. Gene expression data were used for analysis of differential gene expression and analysis of the dynamics of gene expression variations.

**GSE38351:** Smiljanovic et al. generated profiles of human peripheral blood monocytes activated in vivo and stimulated in vitro. There were 15 SLE patients ageing from 21-63 years, 14 RA ageing from 67-22 and 12 Healthy donors aging from 20-60 years were included in the cohort. Monocytes from patients with SLE, RA and from healthy donors were used for generating disease-specific gene-expression profiles, where these profiles represent in vivo activation of monocytes.

In addition, monocytes from healthy donors were stimulated in vitro by cytokines: TNFα, IFNα2a and IFNγ. Cytokine-specific gene-expression profiles were generated by comparing stimulated monocytes with unstimulated ones. TNFα, IFNα2a- and IFNγ as cytokine-specific gene-expression profiles were compared with RA and SLE, as disease-specific gene-expression profiles.

**GSE4588:** In this cohort, CD4 T and B cells were sorted by flow cytometry from PBMC of patients with SLE, RA and healthy controls. GeneChip® Human genome U133 Plus 2.0 arrays were hybridized in monoplicates and the genes differentially expressed among the three groups of patients were identified using ANOVA tests with corrections for multiple comparisons.

**GSE46920**: In this cohort, Monocytes from 3 healthy donors were cultured for 6 hours in the presence of 20% serum from three newly diagnosed, untreated SLE patients. Microarray analysis was then performed upon normalizing the gene expression levels of samples incubated with SLE sera to those incubated with autologous serum.

**GSE46923:** In this cohort, Rodriguez-Pla et al. collected samples from 51 SLE patients, including 7males and 44 females. The average age of the patients at the day of sample collection was 15 years (range: 8–19), and the average duration of SLE was 0.69 years (range: 0–2.06 years). The breakdown of the patient ethnicity was: 45% Hispanic, 27% African American, 18% Caucasian, 4% Asian, and 2% unspecified. Thirteen sera from pediatric SLE patients were used for the flow cytometry staining on cultured monocytes, most of them in more than one independent experiment. SLE sera were selected based on high Systemic Lupus Erythematosus Disease Activity Index (SLEDAI), absence of immunosuppressive medication, and absence of high dose prednisone at the moment of blood draw, in addition to absence of intravenous prednisolone bolus administration in the two months previous to the date of blood draw. Patients were all females. Average age was 15.8 years (range: 13– 17). The breakdown of ethnicity was: 61.5% Hispanic, 30.8% African American, and 7.7% Caucasian.

The control population consisted of 21 randomly selected healthy children, including 5 males and 16 females (average age: 12 years; range: 6–22). The ethnic breakdown of the healthy donors was: 42% Caucasian, 29% Hispanic, 19% African American, and 10% Asians. Some of the flow cytometry staining on cultured monocytes were done using monocytes from three adult healthy donors (two males and one female with ages ranging from 31 to 56 years.)

**GSE50772:** Kennedy et al. enrolled all patients met the American College of Rheumatology criteria for SLE. They registered this trial (<u>NCT00962832</u>) on the ClinicalTrials.gov website. For purposes of executing clinical trials with different end points, patients with SLE are characterised predominantly as patients with extrarenal lupus (ERL) or as patients with lupus nephritis (LN). The following cohorts were evaluated: 61 patients with extrarenal lupus (ERL) in the University of Michigan observational cohort, 60 patients with mild ERL enrolled in the rontalizumab Phase I trial,^16^ 135 patients with moderate-severe ERL in the EXPLORER rituximab trial,^17^ 80 patients with moderate-severe lupus nephritis (LN) in the LUNAR rituximab trial^18^ and 238 patients with moderate-severe ERL in the ROSE rontalizumab (anti-IFN-α monoclonal antibody) trial.^19^ Healthy control subjects (n=85) were recruited by the Genentech blood donation programme for research use of blood samples, and were age matched and gender matched to the lupus trial patients.

**GSE51997: Kyogoku et al.** collected cells from Systemic Lupus Erythematosus (SLE) patients as follows: For CD4pos T cells, six patients with SLE (average age: 29.0 +/- 7.6) and four normal healthy donors (ND; 24.8 +/- 0.5) were recruited. For CD16neg monocytes, four patients with SLE (26.5 +/- 1.7) and four NDs (24.8 +/- 0.5) were recruited. For CD16pos monocytes, four patients with SLE (26.5 +/- 1.7) and three NDs (24.7 +/- 0.6) were recruited. All patients and NDs were female. The same NDs were examined before and after immunisation with yellow fever vaccine (YFV). Collection of cells from yellow-fever vaccinated individuals: ND were immunised with a vaccine against the wild-type YF virus, which is a single-stranded RNA virus without adjuvants. This vaccine consists of a live but attenuated strain of the yellow fever virus (YFV-17D). Based on its vaccination-associated clinical and serological manifestations, this immunisation can be regarded as a real viral infection. A total of 50 ml peripheral blood was taken 7 days after immunisation, when sufficient numbers of CD19pos/CD27high plasmablasts were detected by flow cytometry. Cell sorting: A total of 50 ml peripheral blood was collected in Vacutainer heparin tubes and erythrocytes were lysed in EL buffer. Subsequently, granulocytes were depleted using CD15-conjugated microbeads (MACS). The CD15-depleted fraction was stained with a CD14-fluorescein isothiocyanate (FITC) antibody, a CD16-APC-Cy7 antibody, a CD3-Vioblue antibody and a CD4-FITC antibody. Using a FACSAria cell sorter, CD4pos T cells, CD16neg monocytes and CD16pos monocytes were isolated with purities and viabilities of >97%. After sorting, the cells were immediately lysed with RLT buffer and frozen at −70 °C. Total RNA was isolated using an RNeasy mini kit, and quality control was ensured by Bioanalyser measurements.Total RNA was extracted using the RNeasy Mini kit. The integrity and amount of isolated RNA was assessed for each sample using an Agilent 2100 Bioanalyzer and a NanoDrop ND-1000 spectrophotometer. Biotinylated complementary RNA (cRNA) was synthesized from 100 ng total RNA, using reagents as recommended in the technical manual from Affymetrix. Fifteen micrograms of fragmented cRNA of each sample were hybridized to HG-U133 plus 2.0 arrays. Hybridization was performed according to procedure 2 as described in the technical manual. Finally, the arrays were scanned with a GeneChip Scanner 3000 using the GCOS software. All relevant GCOS data of quality checked microarrays were analyzed with High Performance Chip Data Analysis (HPCDA, unpublished), using the BioRetis database (www.bioretis-analysis.de), as described and validated previously.

**GSE52471:** Jabbari et al. profiled the transcriptome of Discoid lupus erythematosus (DLE) skin in order to identify signaling pathways and cellular signatures that may be targeted for treatment purposes. Further comparison of the DLE transcriptome with that of psoriasis, a useful reference given our extensive knowledge of molecular pathways in this disease, provided a framework to identify potential therapeutic targets. Although a growing body of data support a role for IL-17 and T helper type 17 (Th17) cells in systemic lupus, we show a relative enrichment of IFN-γ-associated genes without that for IL-17-associated genes in DLE. Extraction of T cells from the skin of DLE patients identified a predominance of IFN-γ-producing Th1 cells and an absence of IL-17-producing Th17 cells, complementing the results from whole-skin transcriptomic analyses. These data therefore support investigations into treatments for DLE that target Th1 cells or the IFN-γ signaling pathway.

Eleven patients with active DLE were enrolled in the study. Punch biopsy and shave biopsy specimens of psoriasis (n=5) and normal (n=3) skin samples were from patients with active moderate to severe disease or healthy subjects, respectively. Additional sample data from prior studies were added.

**GSE55447:** Sharma et al. collected Peripheral blood from 21 African-American (AA) and 21 European-American (EA) SLE patients, 5 AA controls, and 5 EA controls. CD4+ T-cells, CD8+ T-cells, monocytes and B cells were purified by flow sorting. Each cell subset from each subject was run on an Illumina HumanHT-12 V4 expression BeadChip array (n=208 arrays). Differentially expressed genes (DEGs) were determined by comparing cases and controls of the same ancestral background.

**GSE72747:** Ducreux et al. recruited Twenty-eight patients with SLE (aged 18–50 years), according to the ACR criteria for SLE, in a multicentre, randomized, double-blind placebo-controlled, phase I/II staggered dose-escalation trial of IFN-K (ClinicalTrials.gov registry number NCT01058343). Patients were randomized to receive three or four injections of placebo (n = 7) or 30 µg (n = 3), 60 µg (n = 6), 120 µg (n = 6) or 240 µg (n = 6) IFN-K.

Global gene expression studies were performed in serial whole blood samples from SLE patients with a renal BILAG A prior to, 3 months, and 6 months after initiation of conventional immunosuppressive therapy (induction with high-dose corticosteroids, IV cyclophosphamide or oral mycophenolate during the first 3 months, followed by maintenance with moderate- to low-dose corticosteroids, azathioprine or mycophenolate). The expression of IFN-induced genes was analyzed, in comparison to global and renal indices of disease activity.

**GSE78193:** Normal donor blood was incubated with or without IFN-g stimulation to establish an IFN-g gene signature. Twenty-six patients aging between 18-65 years, with mild-to-moderate and stable SLE were administered placebo or a single dose of AMG 811, ranging from 2 mg to 180 mg subcutaneously or 60 mg intravenously. Antimalarial agents, leflunomide, azathioprine, methotrexate, and up to 20 mg/day of prednisone (or equivalent) were permitted as concomitant therapies. A therapeutic anti-IFN--g antibody, and changes in the IFN--g signature in whole blood of these subjects was measured.

Whole blood PAXgene tube samples were collected from all cohorts at baseline, day-1 (pre-dose), and at days 15, 56, and end of study (EOS) after treatment Arrays were hybridized in a Loop design.

## Notes

**Conflict of interest statement**: The authors have declared that no conflict of interest exists.

